# Planning-while-acting: addressing the continuous dynamics of planning and action in a virtually embodied task

**DOI:** 10.1101/2024.11.28.625911

**Authors:** Davide Nuzzi, Paul Cisek, Giovanni Pezzulo

**Author notes:** Corresponding author January 22, 2026.

## Abstract

Everyday tasks, such as selecting routes when driving or preparing meals, require making sequences of embodied decisions, in which planning and action processes are intertwined. In this study, we address how people make sequential embodied decisions, requiring balancing between immediate affordances and long-term utilities of alternative action plans. We designed a novel virtually embodied task in which participants controlled an avatar tasked with “crossing rivers” by jumping across rocks. The task permitted us to assess how participants balanced between immediate jumping affordances (“safe” versus “risky” jumps) and the utility (length) of the ensuing paths to the goal. Behavioral and computational analyses revealed that participants planned ahead their path to the goal rather than simply focusing on the most immediate jumping affordances. Furthermore, spatial and embodied components of the task influenced participants’ decision strategies, as participants’ current direction of movement and momentum influenced their choice between safe and risky jumps. Additionally, participants showed (pre)planning before making the first jump, but they continued deliberating during it, with movement speed decreasing at decision points and when approaching them. Computational modeling indicates that farsighted participants who assigned greater weight to the utility of future jumps showed a better performance, highlighting the usefulness of planning in embodied settings. Finally, analysing participants’ performance across trials indicates that during the experiment, they become faster in moving and deciding but they do not change their overall strategy. Our findings underscore the importance of studying decision-making and planning in ecologically valid, embodied settings, providing new insights into the interplay between action and cognition in real-world planning-while-acting scenarios.

**Significance statement:** Everyday activities, such as navigating a crowded street or assembling objects, often involve continuous interactions between planning and action. Yet, it is unclear how people balance opportunities for action (affordances) with long-term planning in embodied settings. In a novel virtually embodied task, in which people have to “cross rivers” by making a series of jumps across rocks, we found that they plan ahead their path to the goal rather than simply focusing on the immediate jumping affordances - and they continue deliberating during movement. Crucially, spatial and embodied constraints, such as movement direction and momentum, influenced decision-making, challenging traditional models that treat decision and action as distinct stages and highlighting the importance of studying decisions in ecologically valid, embodied settings.

## 1 Introduction

Most everyday tasks, such as selecting a path in a crowded street, preparing a table or a backpack for a trekking, require sequences of embodied decisions, in which planning and action processes unfold simultaneously. Traditional studies of decision-making and planning in psychology and (neuro)economics focused instead on static, non-embodied scenarios, such as abstract problem-solving or choices between predefined offers, in which action requirements are minimal. Therefore, it is unclear how individuals plan ahead and how they balance the immediate affordances and the longer-term utility of different courses of actions in embodied settings.

Recent work addressed some of these limitations by focusing on simple embodied decisions and asked how people select between different courses of action, by weighting immediate affordances – or opportunities for action – and utilities (1; 2; 3; 4; 5; 6; 7; 8). However, most of the above studies focused on immediate actions and affordances, such as selecting between reaching, stepping, or pressing actions, or choosing between following a current target or switching to a new one, without addressing *sequential* decisions and planning (9; 10; 11; 12; 13; 14; 15; 16; 17; 18; 19; 20; 21). Some studies (10; 14; 15; 21) asked participants to make embodied decisions while they were already acting. For example, when participants continuously tracked a moving target and were occasionally presented with a new target to which they could freely choose to switch, their decisions were influenced by geometric factors, such as the distance and alignment to the new target (10) as well as biomechanical costs (21). In another study, participants who were already walking had to decide on the fly which lateral (left vs. right) target they would go to, when the two targets were associated with different rewards. This study revealed that current stepping behavior influences the choice of the next target, even if this implied obtaining less reward, therefore highlighting that action dynamics can influence value-based decisions. Still another study used a similar walking paradigm and additionally manipulated the turning cost to reach the lateral targets. It reported that the dynamically changing motor costs significantly influenced the choice of the target (15). While these studies (and others) highlighted that geometric and embodied aspects influence decisions between immediate actions, they did not address the role of such factors in sequential choices between alternative action plans. Therefore, they could not address how people balance immediate affordances versus long-term goals. Likewise, studies investigating decision-making and affordance competition in sports highlighted the importance of embodied strategies for successful performance, but rarely addressed sequential decision settings (22; 23; 24; 25).

An established line of research by Rosenbaum and collaborators addressed more directly how people plan and control sequential physical actions, revealing various interesting findings. An important finding is that when participants had to select between sequences of motor responses, they adopted a hierarchical movement planning strategy and prepared for forthcoming sequence choices by preactivating responses shared by the alternative sequences (26). Another intriguing finding is that, when performing sequences of movements such as picking a bucket and carrying it to the alley’s end, and additionally given the choice between a bucket that was closer to the start position and one that was close to the end, participants selected the former, even if it implied more effort to carry it farther than the other bucket. This finding is indicative of *pre-crastination* (27; 28; 29), or the tendency to get things done as soon as possible, even if that involves extra effort. However, a subsequent study showed that as the costs to reach and carry the bucket increased, participants reversed their preference, showing that pre-crastination depends on cost-benefit considerations (30). Another finding is that people are much (about 5 times) faster in choosing between temporally extended behaviors compared to performing them, suggesting that they might not mentally simulate all the alternatives before deciding between them (31). Other studies in this line of research addressed the coordination of reaching and walking movements, revealing that the cost of reaching was much greater than the cost of walking (32; 33). Yet other studies showed that participants showed *end-state comfort* effects both during manual action and during locomotion, when they selected an initial hand (or feet) posture in order to gain comfort at the end of the sequence (34; 35). While important, these studies largely focused on higher-level choices between action plans. The few studies that addressed sequential effects, such as pre-crastination (27; 28; 29) and the end-state comfort effect (34; 35), focused on very short and rather stereotyped sequences of movement, without addressing planning domains requiring people to construct and compare multiple long-term plans de novo. Furthermore, they did not address whether and how deliberation continues while people execute the plan; hence, they could not fully analyze the possible feedback between movement and subsequent decisions. Finally, they did not formalize the ways participants balance immediate and future affordances when making sequential decisions. Likewise, large scale studies of pedestrian (36) and videgame-based navigation (37; 38) addressed high-level (e.g., vector-based) strategies supporting path planning, but did not systematically investigate fine-grained planning dynamics, such as how participants behave at individual decision points.

While the aforementioned studies advanced our understanding of embodied decisions, especially those that exploit immediate affordances, the ways people solve sequential embodied tasks requiring the continuous construction and evaluation of novel action plans, and the balance between those offering immediate affordances versus those having greater longer-term utility, remain incompletely understood. Importantly, the fields of psychology and (neuro)economics have developed solid theoretical and experimental approaches to address sequential decisions by presenting participants with a series of two-alternative forced choices between offers, such as “safe” versus “risky” offers, associated with different probabilities and utilities, while allowing them to plan multiple steps ahead (39; 40; 41). However, in these fields, sequential decisions are studied in non-embodied settings, in which the movements required to make the first choice are irrelevant for the subsequent choices. An appealing possibility is therefore adopting the same methodology, based upon a series of choices between offers, but translating the classical choice factors – like probabilities and utilities – into an embodied setting, in which decisions and actions are intertwined, i.e., a planning-while-acting setting (23).

Adapting the logic of non-embodied economics sequential tasks allows us to generalize find-ings—and explore differences with—embodied settings. For example, by studying whether and how the factors inherent in sequential decisions made during real physical behavior, such as present and future affordances and the concurrent planning and action dynamics, influence participants’ decisions above and beyond the factors traditionally considered, such as the economic value of the alternative plans. It allows us to address and formalize novel, previously unexplored key aspects of sequential embodied decision-making. For example, in sequential embodied settings, do people prefer plans whose immediate affordances for movement or long-term utility are greater? Do embodied factors of the situation, such as the spatial configuration of the environment, the participants’ direction of movement at decision points, and the cost of changing trajectory, impact sequential choice dynamics, and if so, how? Do people preplan action sequences before initiating movements? Does the planning phase end at movement initiation or continue during movement, i.e., do people simultaneously plan and act?

Here, we address the above questions by designing a novel experiment in which participants control an avatar tasked with “crossing rivers” by jumping across rocks. Crucially, the task requires making a series of jumping choices in a “virtually embodied” setting, in which the movements of the avatar follow coherent spatial and physical rules. Adopting the same sequential structure as non-embodied experiments in psychology and (neuro)economics, we present participants with multiple sequential choices between “safe” versus “risky” jumps, which now incorporate embodied aspects; namely, they correspond to closer-and-bigger versus farther-and-smaller rocks, and between paths having different utilities (here, related to the path length to the goal). Crucially, this design makes choices and movements interdependent: each jump changes the direction and velocity of the avatar and the geometry of the problem. This allows us to study whether and to what extent these factors, ignored in non-embodied settings, influence participants’ sequential decisions.

In sum, by analyzing participants’ behavioral choices and formalizing their strategies using computational models, we aim to reveal how individuals balance affordance and utility in embodied settings, how they form sequential plans, and how their choice strategies are influenced by spatial and embodied aspects of the task. Embodied theories of decision-making predict that participants will be sensitive to both present and future affordances when selecting amongst alternative paths (3). Furthermore, we expect spatial and embodied characteristics of the situation, such as the participants’ current direction of movement, to affect the paths they select at decision points, above and beyond the economic utility of the paths, as predicted by classical, non-embodied theories.

## 2 Methods

### 2.1 Participants

We recruited participants via the Prolific platform, indicating native English and age between 18 and 40 years as eligibility criteria. We only included in the analysis participants who completed all levels of the game, resulting in a sample of 40 participants (19 males and 21 females), with a median age of 30 years, while 5 participants were excluded for not finishing the experiment. All participants gave informed consent online before starting the experiment. All our procedures were approved by the CNR Ethics Committee.

### 2.2 Experimental setup

In this study, we develop a novel 3D video game using Unity (https://unity.com/). The game requires participants to control an avatar (an anthropomorphic frog) to cross rivers, by jumping from rock to rock until a goal platform indicated by a red flag (Figure 1, see also Supplementary Videos S1 and S2). Participants use the mouse to adjust the camera view, the ‘W’ key to move forward, and the spacebar to jump. If the avatar falls into the water, the trial restarts from the last rock on which it stood, after a 2 second animation showing the avatar being transported back. The avatar is 1.75 units tall. The game’s world units are arbitrary, meaning that the relative proportions between them are what matter, rather than their absolute values. The gravity is simulated realistically, with a constant gravitational acceleration of *−*15 units*/*s^2^. Collisions between the character and the platforms, as well as with the water, are handled via an invisible capsule-shaped collider attached to the avatar. A non-realistic simplification was introduced, allowing the character to move forward and rotate even while airborne. However, no additional platforming tricks, such as “coyote time”, were implemented to artificially facilitate movement. The forward movement speed is 5 units per second when the ‘W’ key is pressed. Acceleration from rest to this speed, as well as deceleration when ‘W’ is released, is nearly instantaneous. The vertical speed imparted when pressing ‘SPACE’ to jump is calibrated so that the avatar reaches a maximum height of 1.2 units during a jump. This means the initial vertical speed at takeoff is 6 units per second, the time required to reach the peak is 0.4 seconds, and the total duration of a jump is 0.8 seconds. Keeping SPACE pressed for longer does not alter the height of the jump. If the player holds ‘W’ for the entire duration of the jump, the maximum horizontal distance that can be covered is 4 units. Finally, the avatar’s orientation (i.e., rotation around the vertical axis) is fully controlled by mouse movements. This ensures that the direction the avatar is facing always matches the player’s camera view, meaning there is no discrepancy between where the character is looking and what the player sees.

**Figure 1:**
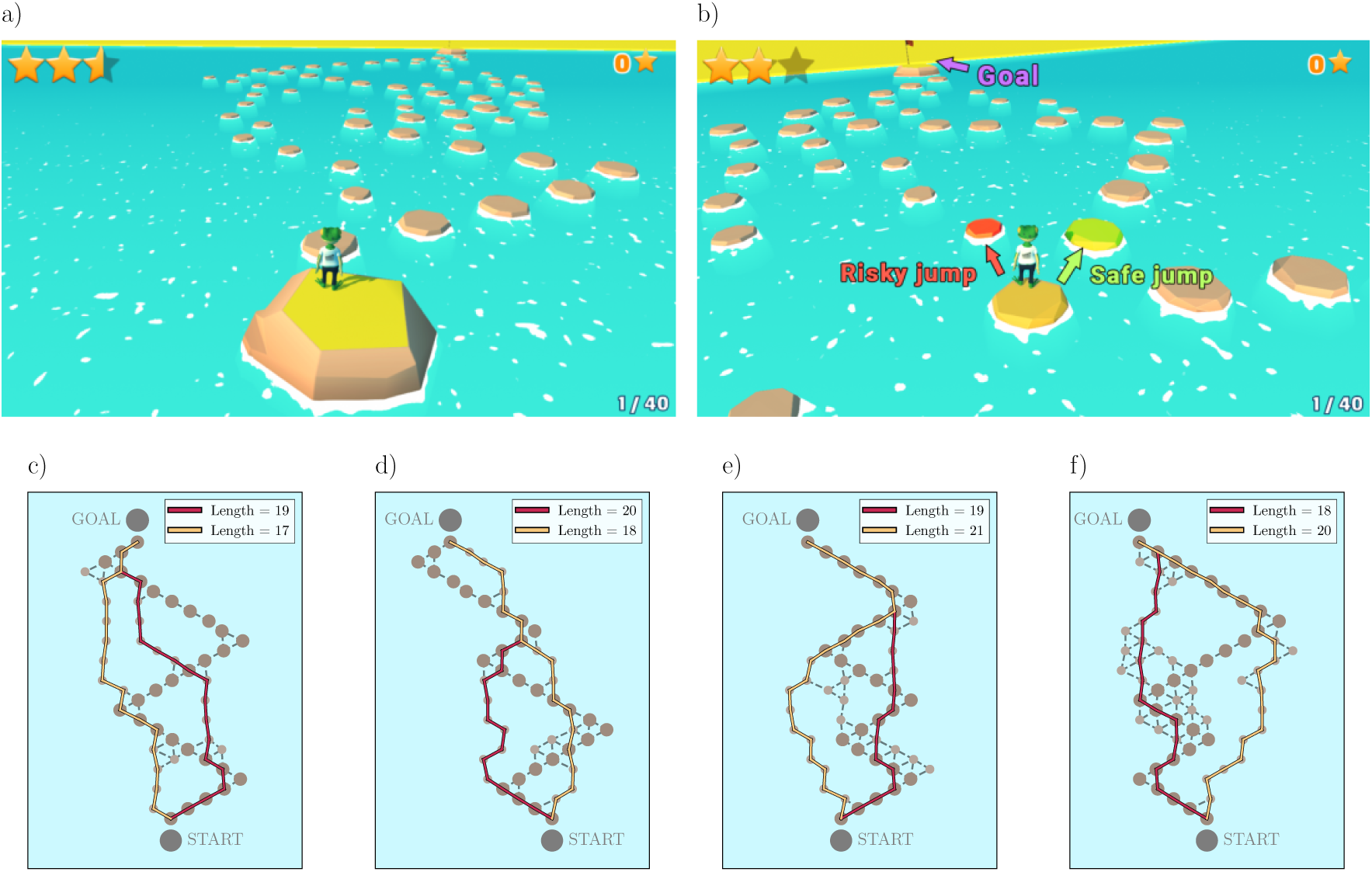
Experimental setup. **a)** A screenshot from the game showing the start of a level. The avatar (frog) is positioned on an elevated platform, providing a complete view of the entire map and the distant goal platform, marked with a red flag. **b)** A screenshot illustrating a decision point where the player must choose between two types of jumps: a “safe” jump to a larger rock or a “risky” jump to a smaller rock. The goal platform is visible in the distance. **c-f)** Examples of the four categories of maps used in the experiment, based on the properties of the first decision point. The first *risky* jump can be on the shorter path to the goal (c,d) or on the longer one (e,f). The direction of the first *risky* jump can have a smaller angle with respect to the goal direction (c,e) or a larger angle (d,f). In each map, the *safe* path is shown in red, and the *risky* paths are shown in yellow. The difference in path length (number of jumps) between the shorter and longer paths is always 2 jumps.

Participants start each level from a starting platform, which provides them with a full view of the map (Figure 1.a). Crossing the river requires jumping across rocks of two types: large rocks, affording a *safe* jump to a larger landing area and small rocks, affording a *risky* jump to a smaller landing area. Platform size and jump distance were not manipulated independently; thus, all safe jumps involved closer and larger platforms, whereas all risky jumps involved farther and smaller platforms, with no intermediate combinations. The large and small rocks colliders have an approximate radius of 0.72 and 0.48 units respectively. The game levels are generated procedurally. They include a *safe* path composed only of large rocks arranged in a zig-zag pattern on a hexagonal grid from the starting platform to the goal platform. Furthermore, they include multiple *risky*paths composed of smaller rocks, which branch off the *safe* path at random points. The *risky* paths move toward the goal by selecting one of three directions on the hexagonal grid: forward left, forward, or forward right. This design creates multiple decision points, between safe or risky jumps and paths. Notably, for each problem, there is a single shortest path, which could be either the *safe* path or a *risky* path. Regardless of which one is shorter, the difference in distance between the shortest path and the other is always two jumps. We fixed the distance between points on the hexagonal grid to 3.9 units, preventing participants from jumping to non-adjacent rocks. Furthermore, we applied a positional jitter to the rocks after the initial map generation, to create a more naturalistic arrangement. Specifically, each platform was moved in a random directions by 0.4 units. We made sure that the randomization was the same for all players and that, after the random jitter, the border-to-border distance between adjacent platforms (w.r.t. the hexagonal grid) was always less than the maximum jump distance of the avatar (4 units).

Our experiment used 40 game levels and a 2 x 2 design. The two factors reflect the decision to be made at the first decision point between a *safe* and a *risky* jump (Figure 1.b). The first factor reflects whether the the shortest path to the goal (measured as the minimum number of jumps) started with the *safe* or the *risky* jump. The second factor is the angle between the vector connecting the decision point to the chosen rock and the vector connecting the decision point to the goal, indicating how aligned the jump and the direction of the goal are. As a single jump is insufficient to indicate the path’s overall direction, we computed an average angular deviation from the goal direction by considering the initial segments of each path. Specifically, at each step along the path, we calculated the angle between the current movement direction and the straight-line direction to the goal, and averaged these values using exponentially decaying weights to prioritize earlier steps. Note that path length is typically a better proxy for performance than the angle to the goal, as fewer jumps generally require less time. However, the angle to the goal provides an immediate, visually perceivable cue, whereas estimating path length is more challenging – especially considering that paths often involve more than 20 jumps. By considering both factors, we aimed to assess how participants balance efficiency and perceptual availability. Based on these two factors, we created four groups of ten levels each, for a total of 40 levels (Figure 1.c-f). We procedurally generated multiple levels for each factor and manually selected 40, ensuring that each selected level had at least 5 decision points.

Participants are informed that they have a maximum of 80 seconds to complete each level and that they are awarded from one to three stars depending on problem completion time (or zero stars if they fail to complete the problem within the time limit). The total score, i.e. the total number of collected stars, was always visible in the top-right corner of the screen. Before the experiment, participants completed three tutorial levels to familiarize with the controls and game mechanics.

### 2.3 Data analysis methods

We used mixed-effects models to account for both fixed effects of experimental factors and random effects associated with individual variability. We employed the *pymer4* package in Python, using the *Lmer* function for both generalized linear mixed models (GLMMs) and linear mixed models (LMMs), depending on the dependent variable.

In the case of binary outcome variables (e.g., probabilities), we used GLMMs with a logit link function, whereas in the case of continuous variables we used LMMs without a logit link function. In both cases, we included in these models random intercepts for each participant, to capture participant-level variability.

For each analysis, we compared models with or without interaction terms between factors, using Bayesian Information Criterion (BIC) and we retained the model having the lower BIC score, indicating a better fit. Following the model fitting procedure, we conducted post-hoc analyses when we found main effects or interactions to be significant. We carried out these post-hoc pairwise comparisons using the *post hoc* function in *pymer4*, with Tukey’s Honest Significant Difference (HSD) correction applied to adjust for multiple comparisons. This procedure ensured that we identified specific differences between conditions while controlling for Type I errors.

We set the statistical significance at *p <* 0.05 for all analyses and we indicated in the boxplots all the significant differences from the post-hoc tests.

## 3 Results

We first considered whether participants were able to solve the game effectively, by looking at their general performance metrics (reported below as mean *±* standard deviation) across the experiment. A total of 26 out of 40 participants completed all trials within the time limit, while others failed to complete one or more levels before the timer expired. On average, participants successfully completed 36.05 *±* 8.46 trials, making 30036 successful jumps and 2583 failed jumps (on average, 21.30 *±* 2.97 jumps per level). The total time they spent actively playing the levels (excluding tutorials and transitions) was 31.47 *±* 14.63 minutes.

Participants demonstrated a high level of accuracy in the task, with an overall jump success rate of 93% *±* 5%, falling into the water 1.61 *±* 2.02 times per level on average. This indicates that errors were relatively rare, and performance was generally robust across conditions. Performance varied across individuals, with participants earning an average of 76.90 *±* 30.90 stars across all levels, out of the 120 possible stars if they completed each level perfectly.

### 3.1 Participants rely on both path length and angle to goal to decide between *safe* or *risky* jumps

We analyzed the probability of making the first *risky* jump (i.e., jumping onto the small rock at the first decision point) in relation to our two experimental factors: namely, the length of the path to the goal starting with the *risky* jump and the angle between the jump direction and the goal. We computed the probabilities per participant across all levels corresponding to the four specific combinations of these factors (Figure 2.a). Participants select the *risky* jump more often when both the path starting with it is shorter and the angle to the goal is smaller; and less often when the path is longer and the angle is larger. When either the path is shorter with a larger angle or longer with a smaller angle, participants selected the *risky* jump with intermediate probabilities.

**Figure 2:**
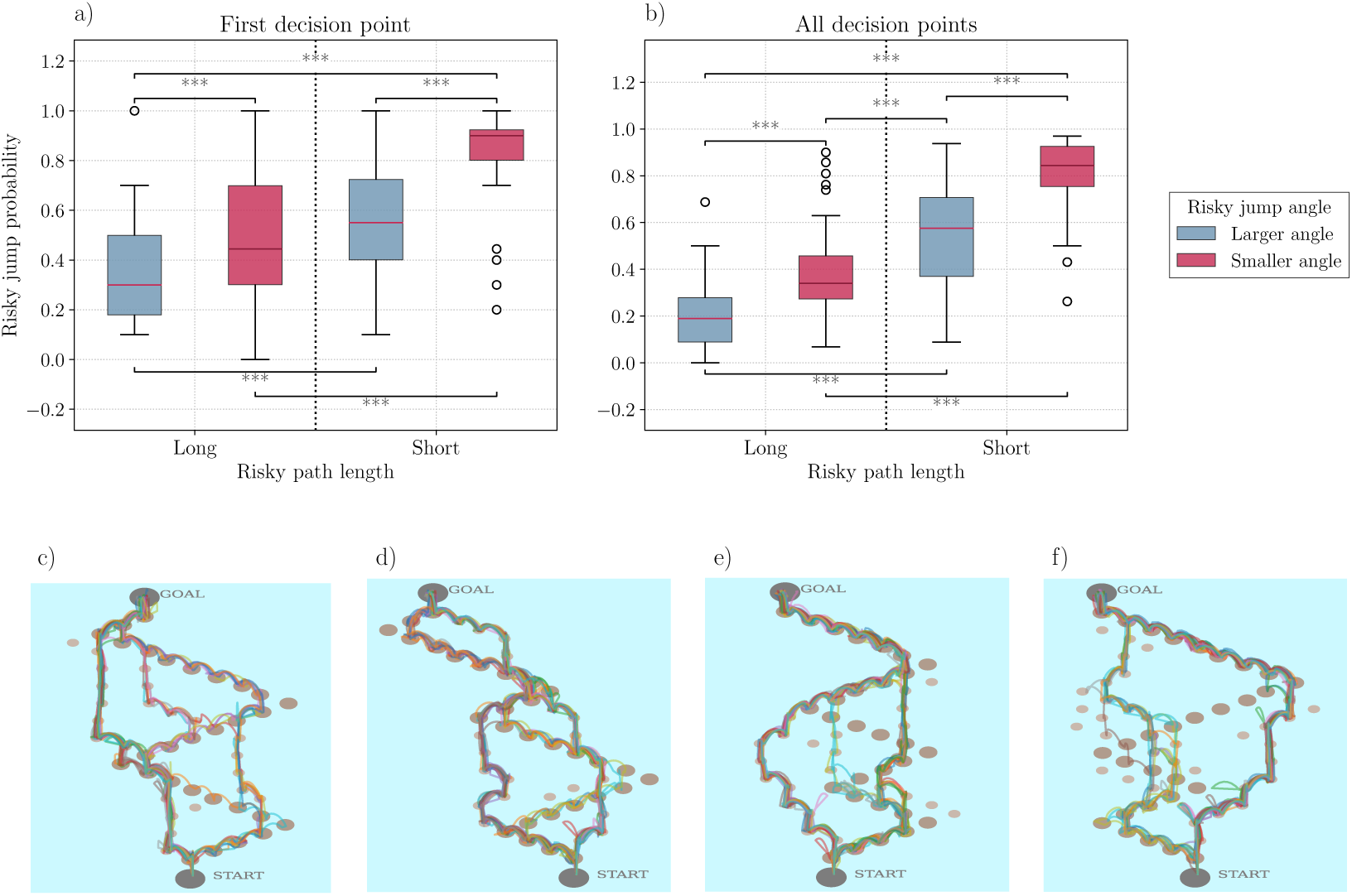
Both path length and angle to goal influence the probability of making a *risky* jump. **a)** Effect of path length and angle on the probability of making the *risky* jump, at the first decision point. Participants show a greater preference for the *risky* jump when the path is shorter and the angle is smaller, and a smaller preference when the path is longer and the angle is larger. There is no significant difference between the two intermediate conditions (shorter path with a larger angle vs. longer path with a smaller angle). **b)** Effect of path length and angle to the goal on the probability of making a *risky* jump, across all decision points. Both path length and angle significantly influence the probability of making a *risky* jump, with path length having a stronger effect than angle to the goal. **c-f)** Trajectories of all 40 participants across the same levels of Figure 1, showcasing the diversity of paths taken at each decision point. Each colored path corresponds to the trajectory of a single participant. Small loops visible in some trajectories indicate points where participants fell into the water and were automatically transported back to their last platform.

To assess the relative influence of the two experimental factors on the probability of taking a *risky* jump, we fitted a generalized linear mixed-effects model (GLMM). We considered a model that includes an interaction term between the factors, as this resulted in a lower BIC than a model without interaction (Table 1). Both path length and angle to the goal significantly affect the probability of choosing the *risky* jump (*p <* 0.001). However, the coefficient for path length is larger, suggesting a stronger influence of path length compared to angle to the goal. Furthermore, the results reveal a significant interaction (*p <* 0.001) between the two analysed factors, highlighting that the probability of risky jumps further increases when the path is shorter and the angle to the goal is smaller. We compared all pairs of conditions using Post-hoc tests (Figure 2.a and Table S3). All the differences are significant (*p <* 0.001), except for the two intermediate conditions (shorter path length, larger angle vs longer path length, smaller angle), indicating a lack of clear dominance of either factor.

**Table 1:** GLMM results for the *risky* jump probability at the first decision point. Both path length and angle to the goal significantly influence the probability of choosing the *risky* jump. The larger *β* coefficient for path length compared to angle to the goal suggests a stronger effect of path length. The interaction between the factors is significant, too.

|  | Fixed effects |  |  |  |  | Random effects |
| --- | --- | --- | --- | --- | --- | --- |
|  | Estimate | Std Error | [0.025 | 0.095] | p-value | Variance |
| Intercept | -0.787 | 0.176 | -1.132 | -0.441 | < **0.001** | 0.719 |
| Risky Jump Shorter Path | 1.082 | 0.158 | 0.773 | 1.391 | < **0.001** |  |
| Risky Jump Smaller Angle | 0.721 | 0.161 | 0.405 | 1.037 | < **0.001** |  |
| Interaction | 0.852 | 0.242 | 0.377 | 1.326 | < **0.001** |  |

These results suggest that participants consider both path length and angle to the goal when deciding which rock to jump onto at the first decision point. Moreover, the impact of path length on the choice is greater than angle to the goal (as quantified by the coefficient estimate), despite the small difference (two jumps) between the lengths of *risky* and *safe* paths and the fact that the angle to the goal is more readily available perceptually.

We extended the analysis to include all decision points, not just the first one. A decision point was defined as a condition where participants faced a forced choice between a *risky* and a *safe* jump. We excluded decisions points with more than two options or where both options were either *risky* or *safe*.

For each decision point, we calculated the length of the path and the angle to the goal for both options and considered whether the *risky* jump was associated with the shorter path or smaller angle than the *safe* jump, forming the same four categories as in the analysis of the first decision point. We then computed the probability of choosing the *risky* jump for each participant across these categories (Figure 2.b). As with the first decision point, we fitted a GLMM to the data (Table 2). This time, we considered the model without the interaction term, as it resulted in a smaller BIC. The results of the GLMM confirm that both path length and angle to the goal are significant predictors of *risky* jump probability (*p <* 0.001). This indicates that participants consistently rely on both factors when making decisions across all decision points, not just at the first decision point. Post-hoc comparisons reveal significant differences across all conditions (*p <* 0.001, Table S4), with path length again exerting a stronger influence than angle, as evidenced by both the coefficient sizes and the pairwise tests.

**Table 2:** GLMM results for *risky* jump probability, across all decision points. Both path length and angle to the goal significantly affect the probability of selecting a *risky* jump. Path length has a stronger effect than angle to the goal, as shown by the size of the coefficients and the results of pairwise post-hoc tests.

|  | Fixed effects |  |  |  |  | Random effects |
| --- | --- | --- | --- | --- | --- | --- |
|  | Estimate | Std Error | [0.025 | 0.095] | p-value | Variance |
| Intercept | -1.610 | 0.133 | -1.870 | -1.349 | < **0.001** | 0.478 |
| Risky Jump Shorter Path | 1.904 | 0.072 | 1.762 | 2.046 | < **0.001** |  |
| Risky Jump Smaller Angle | 1.230 | 0.075 | 1.084 | 1.377 | < **0.001** |  |

### 3.2 Current travel direction influences subsequent jump choices

We next tested the hypothesis that participants’ choice of the next rock was influenced by spatial and embodied aspects of the task – specifically, by participants’ current travel direction at the moment they made the choice.

To test this hypothesis, we investigated whether the direction of travel before reaching a decision point affects jump choice probability. We analysed 3748 individual decisions. For each decision point, we defined two options based on the angle relative to the previous trajectory: “Current direction” (option more aligned with previous direction) and “New direction” (option less aligned with previous direction). To ensure clearly distinct categories, we only included decision points where the angular difference between options exceeded 20^◦^. A schematic illustration of this is shown in Figure 3.a. A statistical comparison with the Mann-whitney U-test reveals that the probability of choosing a specific jump is significantly higher for jumps in the “Current direction” than for jumps in the “New direction” (*p <* 0.001). See the leftmost panel of Figure 3.b.

**Figure 3:**
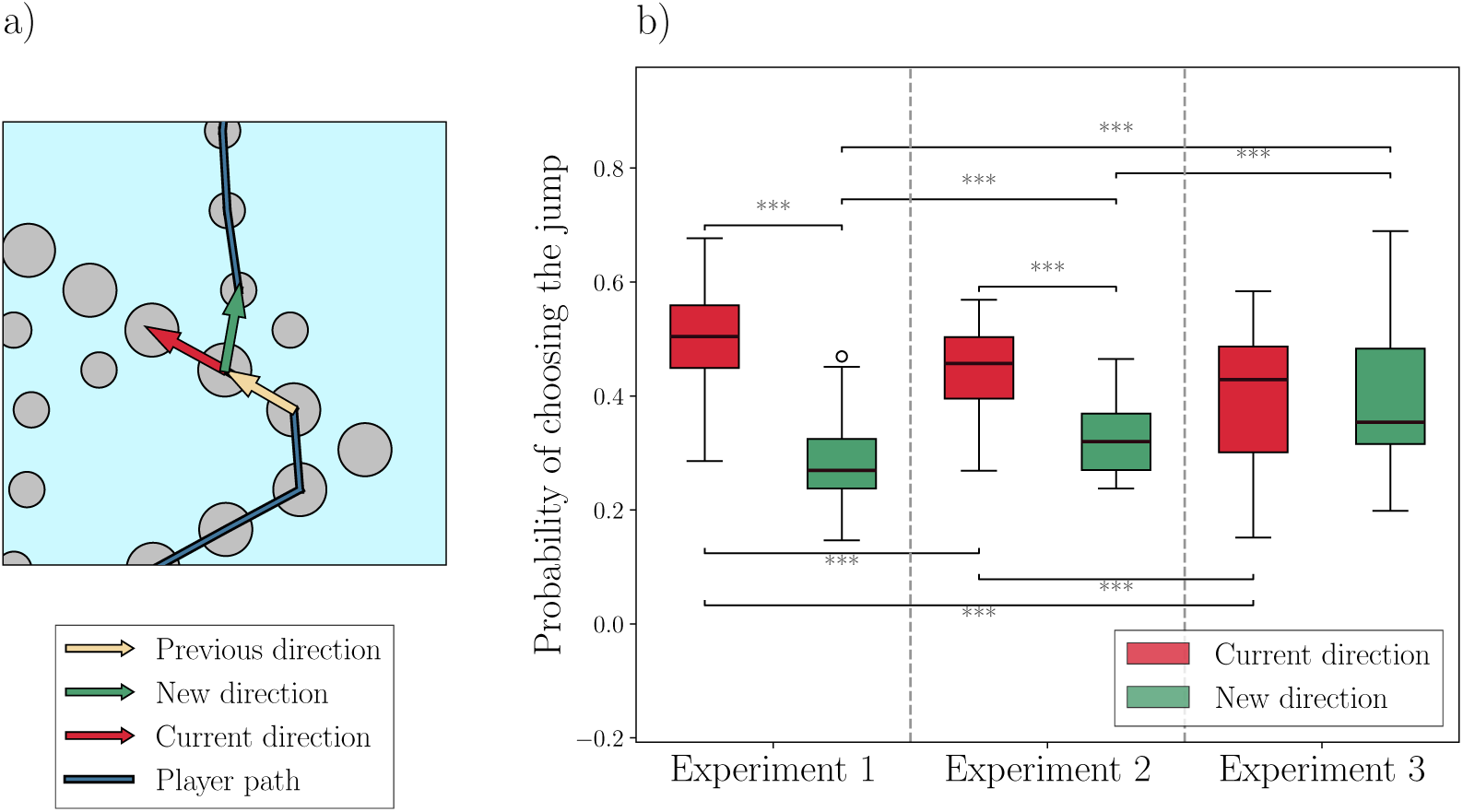
Influence of trajectory direction on jump probabilities. **a)** Illustration of previous, current and new directions at an example decision point. In this example the previous and current directions are almost identical, whereas the new direction, towards a risky rock, is at an angle of approximately 60^◦^ with respect to the previous direction. **b)** Boxplot showing the probability of performing a jump as a function of the direction of the jump, relative to the previous direction, for each of the three experiment. Significant differences are observed across all comparisons (*p <* 0.001) except for the comparison “Current Direction” / “New Direction” in Experiment 2.

We next investigated whether participants’ preference for maintaining the same travel direction depends on spatial factors (e.g., a tendency to plan approximately straight paths), embodied factors (e.g., the influence of physical momentum), temporal or motor costs (e.g., time and/or effort required to change travel direction), or a combination of the above factors. For this, we conducted two control experiments, using the same setup as the main experiment (called Experiment 1 in this analysis) but introduced constraints in the way participants could move the avatar. Experiment 2 (*N* = 30 new participants) introduced a constraint where avatar movements were blocked for 1 second on each rock; the avatars could be rotated but could not jump to another rock. This manipulation breaks the sensorimotor contingency of Experiment 1, removing the effect of physical momentum but not of spatial constraints or motor cost on jumping decisions. Experiment 3 (*N* = 27 new participants) blocked jumps for 1 second like Experiment 2, but additionally rotated the avatars at each decision point, such that they faced the option associated with the “New direction” rather than the “Current direction”. This manipulation introduces a “reverse” effect of motor cost on the choice, such that the higher motor cost is required to select the jumping option associated with the “Current direction”.

We analyzed 8851 individual decisions (comprising Experiments 1, 2 and 3). We modeled the probability of selecting a specific jump using a Generalized Linear Model (GLM) without a global intercept to estimate the effects for all experimental factors directly, including interactions between factors.

The analysis (Table 3) revealed a significant, negative coefficient for the “New direction” factor (Estimate: *−*1.282, *p <* 0.001), indicating a general preference for maintaining the current travel direction, independent of the other factors. The effect of “New direction” interacted significantly with the experiments. The interaction term for Experiment 2 was positive (Estimate: 0.591, *p <* 0.001), indicating that the negative bias against the “New direction” was significantly attenuated compared to the baseline, suggesting a modulatory effect of physical momentum on the choice. The interaction term for Experiment 3 was also positive and larger in magnitude (Estimate: 1.281, *p <* 0.001), indicating that the negative bias was effectively canceled (*−*1.282 + 1.281 *≈* 0). Importantly, the bias was not reversed in favor of the “New direction” as one might have expected if participants prioritized choices associated with smaller motor effort.

**Table 3:** GLM results for jump probability as a function of jump direction, across all decision points in three experiments. All factors show a significant effect except for Experiment 3. The estimate for the factor “New direction” is negative, indicating that choosing a new direction is less common than continuing on the current one, regardless of the experiment.

| Factors | Estimate | Std Error | [0.025 | 0.095] | p-value |
| --- | --- | --- | --- | --- | --- |
| New direction | -1.282 | 0.049 | -1.378 | -1.187 | < **0.001** |
| Experiment 1 | 0.641 | 0.034 | 0.574 | 0.708 | < **0.001** |
| Experiment 2 | 0.346 | 0.041 | 0.266 | 0.425 | < **0.001** |
| Experiment 3 | 0.001 | 0.039 | -0.076 | 0.078 | 0.984 |
| Experiment 2 × New direction | 0.591 | 0.075 | 0.444 | 0.739 | < **0.001** |
| Experiment 3 × New direction | 1.281 | 0.074 | 1.136 | 1.425 | < **0.001** |

Consistent with the above results, the preference for maintaining the current travel direction is evident by the positive coefficient for Experiment 1 (Estimate: 0.641, *p <* 0.001), is still present but attenuated in Experiment 2 (Estimate: 0.346, *p <* 0.001) in which participants were paused, and disappears (log-odds of 50/50) in Experiment 3 (Estimate: 0.001, *p* = 0.984) in which participants were both paused and rotated. Finally, post-hoc comparisons reveal significant differences across all conditions (*p <* 0.001, Table S9) except for the comparison between “Current direction” and “New direction” in Experiment 3, which is not significant. These results are summarized in Figure 3.b.

Summing up, this pattern of results suggests that participants show a preference for following spatially coherent plans, which is attenuated when the effect of physical momentum is removed, and canceled but not reversed when participants are rotated, as would be expected if they simply followed the lower-cost route.

### 3.3 Preplanning time indexes initial decision dynamics

We next examined the relations between the time participants spent preplanning before movement and their subsequent performance and strategy during the levels. We defined *preplanning time* as the time participants spent stationary (without any movement or rotation) on the first rock, before the beginning of movement. If a participant rotated the avatar, the stationary intervals before and after rotation were treated as separate preplanning intervals and summed to compute the total preplanning time on that platform. To assess how preplanning time influenced participants’ performance during the trial, we considered the length of the path they selected. We preferred this measure compared to possible alternatives such as total level score or completion time, since they are confounded by factors such as participants’ skill with the game and are therefore insufficient to isolate planning processes.

We used a linear mixed-effects model (LMM) to analyze the relationship between preplanning time and length of the selected path (Table 4) We found a significant negative effect of preplanning time on path length (*p <* 0.001), showing that participants who spent more time preplanning required fewer jumps to complete the level. A companion, correlation analysis between preplanning time and path length shows a strong negative correlation between the two variables (*R*^2^ = 0.82, Figure 4). Quantitatively, each additional second of preplanning reduces path length by approximately 0.274 jumps.

**Table 4:** Results of the LMM examining the effect of pre-planning time on the length of the selected path. The pre-planning time significantly affects (*p <* 0.001) the length of the path chosen by each participant.

|  | Fixed effects |  |  |  |  | Random effects |
| --- | --- | --- | --- | --- | --- | --- |
|  | Estimate | Std Error | [0.025 | 0.095] | p-value | Variance |
| Intercept | 20.143 | 0.131 | 19.886 | 20.401 | < **0.001** | 0.544 |
| Pre-planning time | -0.274 | 0.047 | -0.366 | -0.183 | < **0.001** |  |

Next, we analyzed how preplanning time was influenced by the same two factors considered in Section 3.1: path length and angle to the goal, both binarized. Specifically, the path length factor indicates whether the shortest path starting from the *risky* jump was shorter or longer than the shortest path starting from the *safe* jump. The angle-to-goal factor compares the alignment between the jump direction and the direction from the current platform to the goal, indicating whether the *risky* jump was more or less aligned with the goal than the *safe* one. We tested the effect of these two factors on preplanning time using a linear mixed-effects model (Table 5). We found no significant difference in preplanning time was observed between levels where the *risky* jump resulted in a longer or shorter path. However, we found that participants spent significantly less time preplanning when the *risky* jump had a smaller angle to the goal (*p* = 0.001). Post-hoc comparisons show participants preplanned faster when the *risky* jump had a smaller compared to a larger angle to the goal (Tables S5). Results of the post-hoc comparisons are shown in Figure 4.b, which displays the marginal effects of the fixed factors after accounting for participant-level variability via random intercepts. These findings suggest that participants required less preplanning when risky jumps were more closely aligned with the goal direction, potentially reflecting a more straightforward or less ambiguous decision.

**Figure 4:**
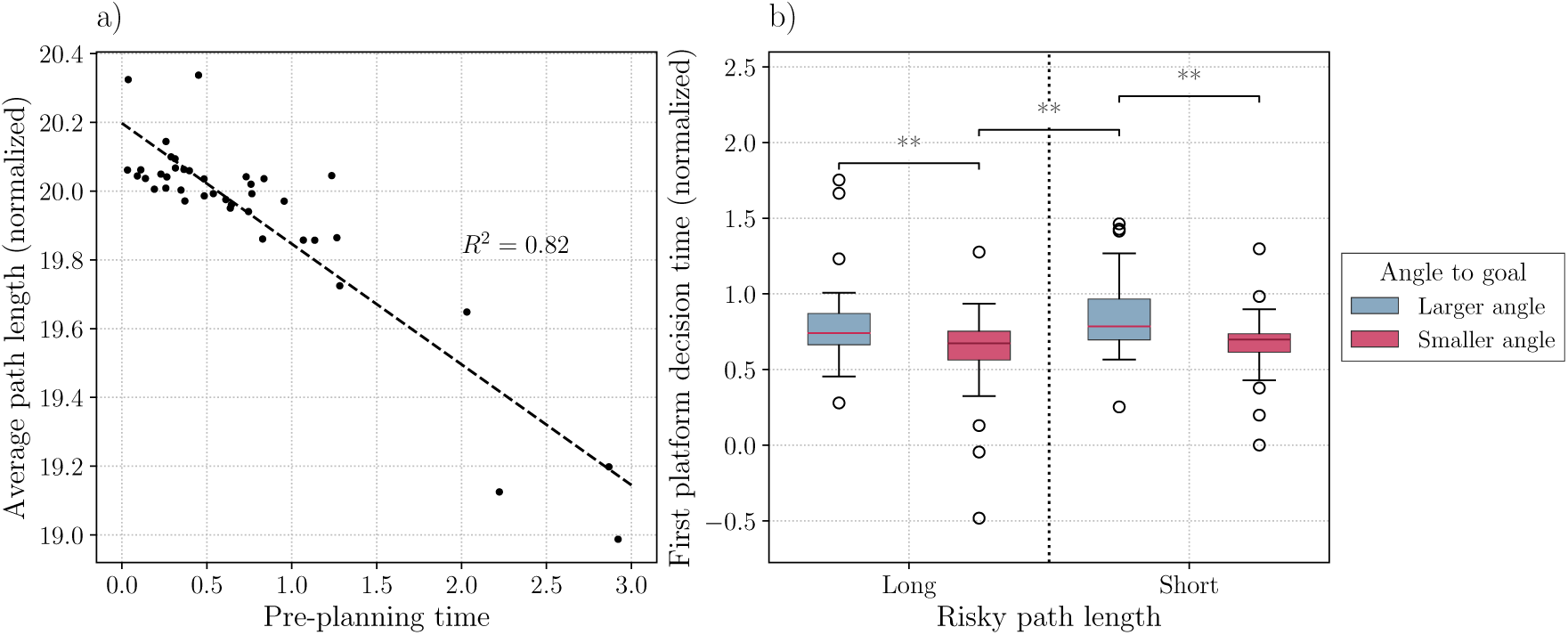
Preplanning time and its effects on path length and decision-making. **a)** Negative correlation between preplanning time (measured in seconds) and average path length (*R*^2^ = 0.82), indicating that participants who spent more time planning required fewer jumps to complete the level. **b)** Boxplot showing decision time as a function of *risky* path length (long/short) and angle to the goal (larger/smaller). Significant post-hoc contrasts reveal that participants pre-plan faster when the *risky* jump has a smaller angle to the goal compared with the safe jump, with no significant difference in pre-planning time based on path length. Time values have been normalized for visualization purposes by removing the per-participant random intercepts from the LMM model. This adjustment removes baseline differences across participants, isolating the effects of the fixed factors.

**Table 5:** Results of the LMM for pre-planning time as a function of *risky* path length and angle to the goal. There is no significant effect of the *risky* path length on the pre-planning time, whereas the angle factor has a significant impact (*p* = 0.001).

|  | Fixed effects |  |  |  |  | Random effects |
| --- | --- | --- | --- | --- | --- | --- |
|  | Estimate | Std Error | [0.025 | 0.095] | p-value | Variance |
| Intercept | 0.792 | 0.116 | 0.564 | 1.019 | < **0.001** | 0.459 |
| Risky Jump Shorter Path | 0.061 | 0.052 | -0.040 | 0.163 | 0.237 |  |
| Risky Jump Smaller Angle | -0.171 | 0.052 | -0.272 | -0.070 | **0.001** |  |

The two analyses of this section highlight distinctive roles of preplanning for selecting the first jump versus the overall path. Despite the fact that preplanning time is not affected by the length of the *risky* path evaluated at the first decision point, it strongly correlates with participants’ ability to select an efficient (shorter) solution to the overall problem, suggesting that during preplanning participants might focus on a global evaluation of the problem rather than only on the first decision.

### 3.4 Planning time during navigation increases at decision points and at platforms preceding them

We next tested the hypothesis that in a continuous task such as crossing the river, planning processes might not only occur at the beginning of a level (i.e., preplanning), but rather continue during the level. To investigate this, we defined *planning time* as the time participants remain stationary on a platform during the solution of a level, without movement or rotation. If a participant rotated the avatar, the stationary intervals before and after rotation were treated as separate planning intervals and summed to compute the total planning time on that platform. We then tested whether planning time increases at decision points and/or at platforms immediately preceding decision points, compared to the other platforms. We categorized platforms into three types: i) *Decision points*, where participants must choose between different risky or safe jumps (as previously described); ii) *Pre-decision points*, platforms immediately preceding a decision point, which are not themselves decision points; iii) *Non-decision points*, all other platforms that are neither decision points nor pre-decision points; see Figure 5.a for a schematic illustration of these platform types in an example level.

**Figure 5:**
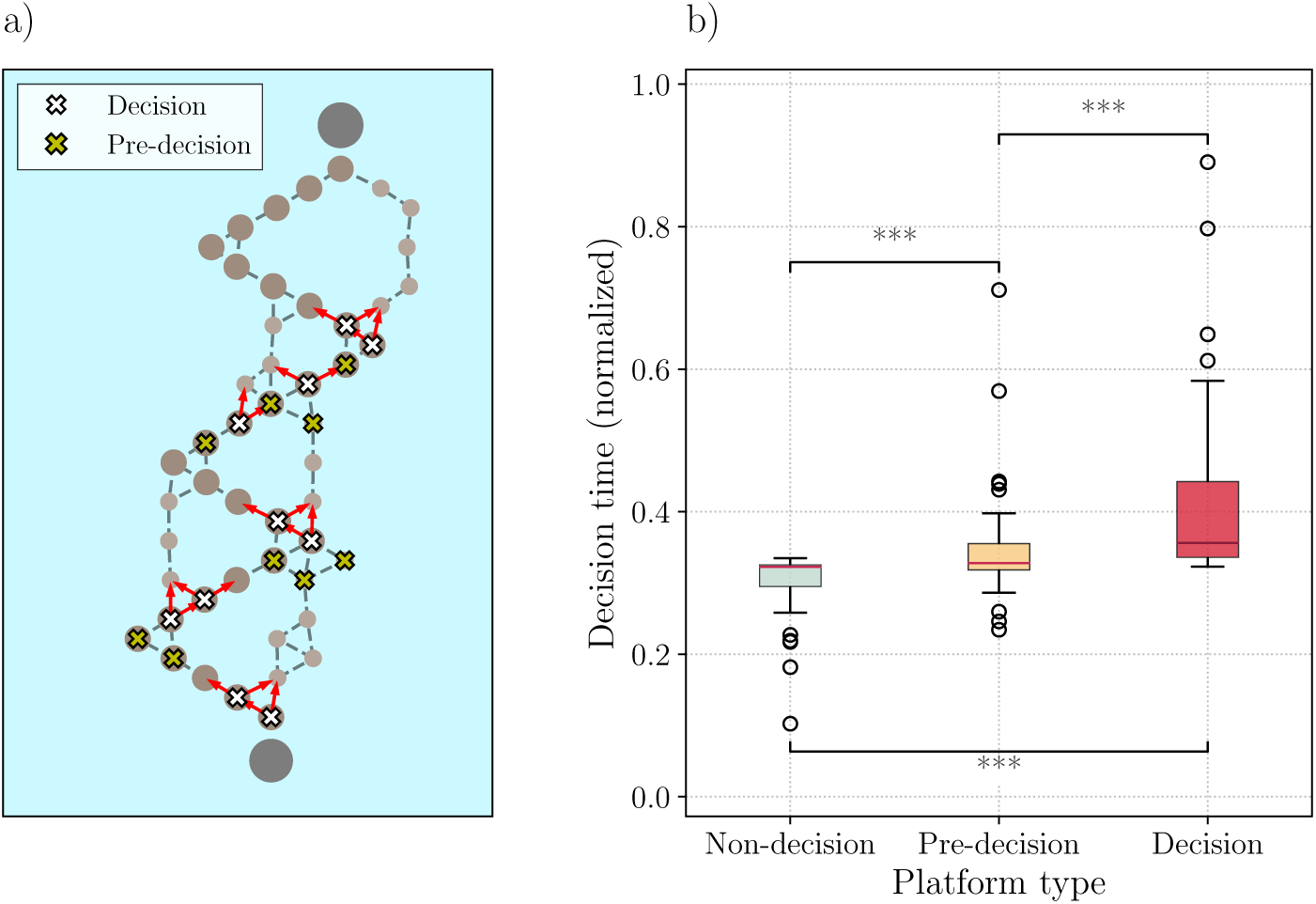
Planning time distribution across different platform types. **a)** Example level showing the distribution of three platform types: decision points, pre-decision points, and non-decision points. Decision points are platforms where participants make choices between risky and safe jumps, pre-decision points are the platforms immediately preceding a decision point, and non-decision points are all other platforms. **b)** Boxplot illustrating the distribution of planning time across the three platform types. Significant differences between platform types are indicated by lines, showing that decision points have the longest planning time, followed by pre-decision points, while non-decision points exhibit the shortest times. Post-hoc comparisons show significant differences (*p <* 0.001) across all three platform types.

We fit a Linear Mixed Model (LMM) to predict planning time based on platform type, considered as a categorical variable with three classes (Table 6 and Figure 5.b). The results show greater planning time for both pre-decision points and decision points (*p <* 0.001) compared to non-decision points, which was taken as the baseline. The coefficient estimates indicate that the increase in planning time is smaller for pre-decision points (*β* = 0.049) than for decision points (*β* = 0.114). See also Table S8 for post-hoc comparisons. These results suggests that planning is a continuous process during crossing the river: it occurs not only before the movement but continues during the task. Furthermore, it is distributed across platforms rather than only occurring at the decision points. Finally, we repeated the analysis using an alternative definition of planning time, which included both the time spent standing still and the time spent rotating on the platform. The results remained consistent: planning time was still significantly longer at decision points compared to pre-decision points (*p <* 0.001).

**Table 6:**
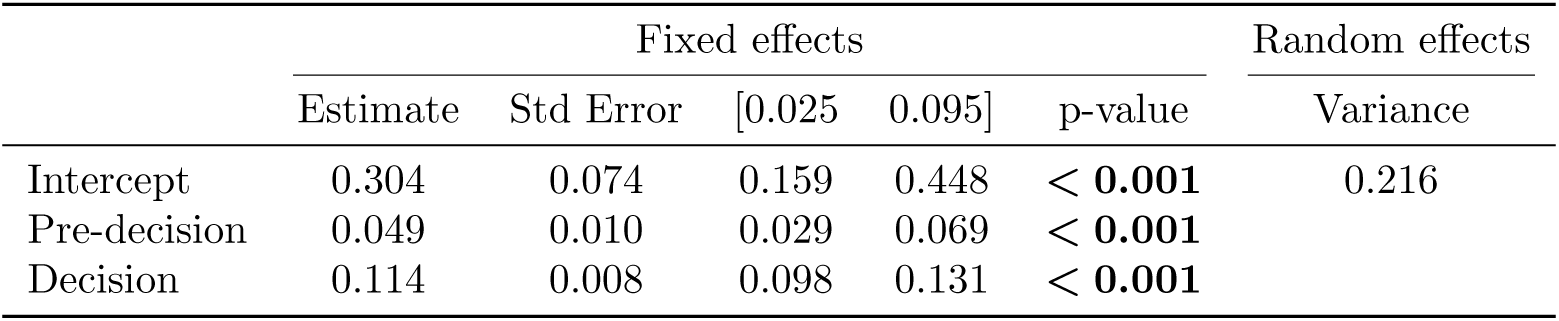
LMM results for planning time as a function of platform type. The table shows that the time spent planning on pre-decision and decision points is significantly different (*p <* 0.001) from the time spent on non-decision platforms, which serve as the baseline.

|  | Fixed effects |  |  |  |  | Random effects |
| --- | --- | --- | --- | --- | --- | --- |
|  | Estimate | Std Error | [0.025 | 0.095] | p-value | Variance |
| Intercept | 0.304 | 0.074 | 0.159 | 0.448 | < **0.001** | 0.216 |
| Pre-decision | 0.049 | 0.010 | 0.029 | 0.069 | < **0.001** |  |
| Decision | 0.114 | 0.008 | 0.098 | 0.131 | < **0.001** |  |

### 3.5 Computational model

We sought to formalize the sequential decision process that participants face in the experiment, using a simple sequential decision model based upon the notion of “expected value” (EV) (42). The model assigns an EV to each jump, defined as the combination of two components: utility (U), which in this case corresponds to a fixed negative cost of −1 per step (as in standard navigation models in reinforcement learning), and affordance (A), which captures the “jumpability” or “landability” of the destination platform. The affordance term replaces the notion of outcome probability in classical economic models (23), modulating the value of an action according to the relative difficulty of executing the jump. Formally, the EV can be expressed as:

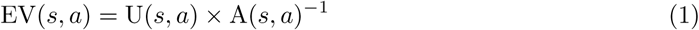

where *s* is the current state of the avatar (the platform it occupies) and *a* is the action (the choice of jumping to one of the neighboring platforms).

This formulation allows actions leading to more accessible platforms to yield less negative values (i.e., closer to zero), aligning with the idea that such jumps are more desirable. Because utility is always negative, dividing by the affordance ensures that higher affordance increases the EV.

To extend the model to sequential jumps, we incorporated a forward-looking aspect, taking into account future jumping choices. For this, we included a discount factor *γ*, which controls how much weight is given to future jumping choices. The revised formula for expected value becomes:

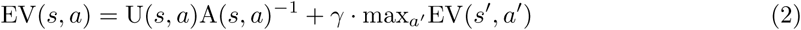

where *s*^′^ = *f* (*s, a*) represents the new state reached after action *a* is taken, and *a*^′^ denotes the available actions from that new state. The discount factor *γ* determines how far-sighted the agent is. With *γ* = 0, the agent only considers the immediate outcome, whereas with *γ* = 1 it gives equal consideration to future decisions. Intermediate values of *γ* represent varying degrees of far-sightedness.

In this model, utility *U* (*s, a*) is defined as a function of path length. If the state *s* is the goal platform, then *U* (*s, a*) = 0, as no further jumps are needed. For all other states, *U* (*s, a*) = *−*1. This negative value ensures that, when affordances are ignored (i.e., when *A*(*s, a*) = 1), the expected value EV(*s, a*) with *γ* = 1 simply corresponds to the number of jumps required to reach the goal, via the shortest path.

Affordance *A*(*s, a*) indexes jumpability and has larger values for easier jumps and smaller values for more challenging jumps. While previous sections explored various types of affordances, here for simplicity we only consider platform distance. While on average greater distance implies a small platform and a *risky* jump, there might be small differences in the distances due to the jitter of rocks – hence considering distance permits a more fine-grained evaluation compared to only considering *risky* and *safe* jumps.

Formally, affordance is defined as:

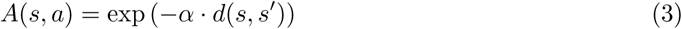

where *d*(*s, s*^′^) is the distance between the current platform and the destination platform after jump *a* and *α* is a coefficient that controls the weight of this factor. To ensure that the model focuses on relative rather than absolute distances, we normalize the affordance values for each platform, such that the maximum value is set to 1 and the other values are scaled accordingly. This scaling allows the model to treat all platforms equally, with only the relative difficulty of jumps influencing the affordance.

Solving this system of equations iteratively, we can compute the EV for each jump, at every decision point. Figure 6 shows landscapes of EVs of an example level, normalized in such a way that greater values (closer to one) correspond to better choices. The figure also illustrates the impact of different values of the model parameters *α* and *γ*. Each row corresponds to a different value of *α*, representing the weight of the affordance term, while each column shows a different value of *γ*, representing the degree of farsightedness. For selected decision points in each level, the figure shows the EV of the two available choices (*risky* and *safe* jumps) as a color gradient on the destination platform, illustrating how the model balances affordance and utility under different conditions.

**Figure 6:**
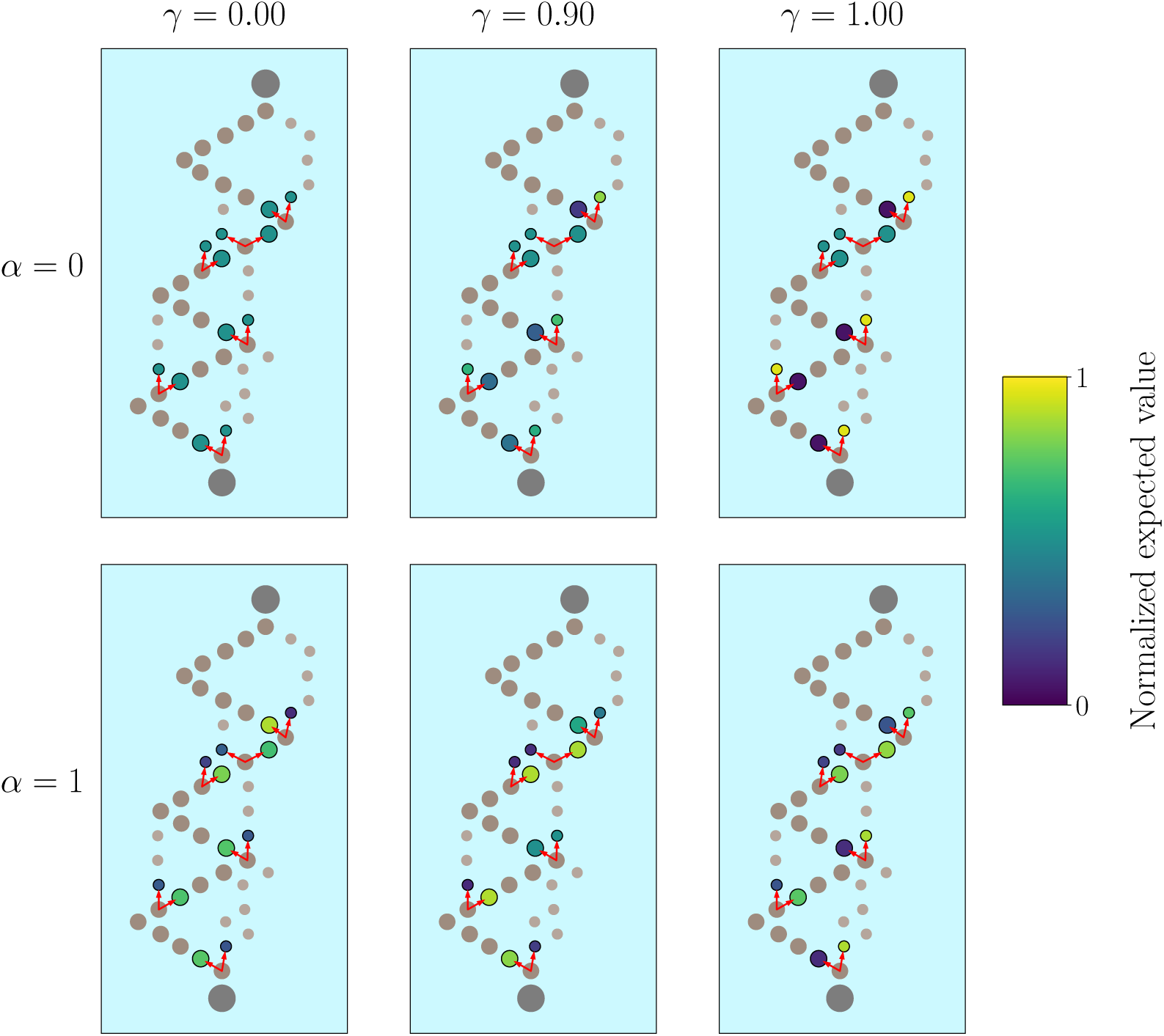
Normalized Expected Value (EV) landscapes for an example problem. The figure shows how normalized EVs change as a function of the parameters *α* (rows) and *γ* (columns) parameters, reflecting affordances and farsightedness, respectively. The color of each platform represents the normalized EV for the corresponding action, with a gradient from 0 (dark blue, representing less favorable actions) to 1 (yellow, indicating more favorable actions). In the top row (*α* = 0), where affordances do not vary with distance, all jumps have the same affordance. For *γ* = 0, the agent evaluates all jumps with the same EV, since it does not consider future jumps. As *γ* increases (moving from left to right), the EV increases for platforms along the shortest path toward the goal, reflecting the influence of planning. In the bottom row (*α* = 1), where the weight of distance-based affordances increases, the model starts to favor safer jumps to closer platforms. This tendency is evident even for *γ* = 0.9, where jumping decisions are also influenced by future jumps. For *γ* = 1, the agent shows a strong preference for the shortest path, while still slightly favoring safe jumps.

To fit the model to participants’ behavior, we convert the expected values into a policy using a softmax function:

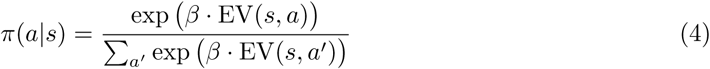

We set *β* = 3, strongly biasing the policy towards the action with the highest EV. For each participant, we measure the negative log-likelihood of their chosen path under the model, using the following equation:

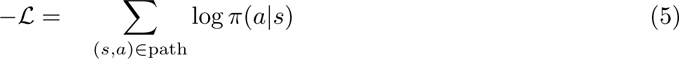

We sum this quantity for each of the 40 levels to get the total log-likelihood. We then minimize the total log-likelihood with respect to the two free parameters, *γ* and *α*, for each participant.

The fitted values of *γ* range from 0.798 to 0.981, indicating that all participants engage in some degree of forward planning while solving the levels. The relatively narrow range of *γ* values can be explained by considering that small variations in this parameter result in large differences in planning behavior, due to its exponential effect on future choices as the number of steps increases. The fitted affordance coefficient *α* reveals two groups of participants. For 15 participants, the values of *α* are equal to zero, suggesting that this group does not prioritize affordance (jump distance) when evaluating the EV of their choices. In other words, these participants adopt a rather uniform strategy, treating all jumps similarly, regardless of distance. By contrast, the *α* of the other participants range between 0 and 0.5, implying that they consider the affordance (distance between platforms) when making jumping choices.

We explored whether the two parameters, farsightedness *γ* and affordance coefficient *α*, correlate with participant performance in the experiment (Figure 7). Our results reveal a significant positive correlation between *γ* and participant performance, with more farsighted participants (those having higher *γ* values) achieving better scores in the task (Pearson *r* = 0.53, *p <* 0.001; Figure 7.a). Furthermore, our results reveal a negative correlation between *α* and the score, with participants giving more weight to affordances (those having higher *α* values) achieving lower scores (Pearson *r* = *−*0.44, *p* = 0.005; Figure 7.b). These results suggest that high-performing participants might pay little attention to the distance between platforms.

**Figure 7:**
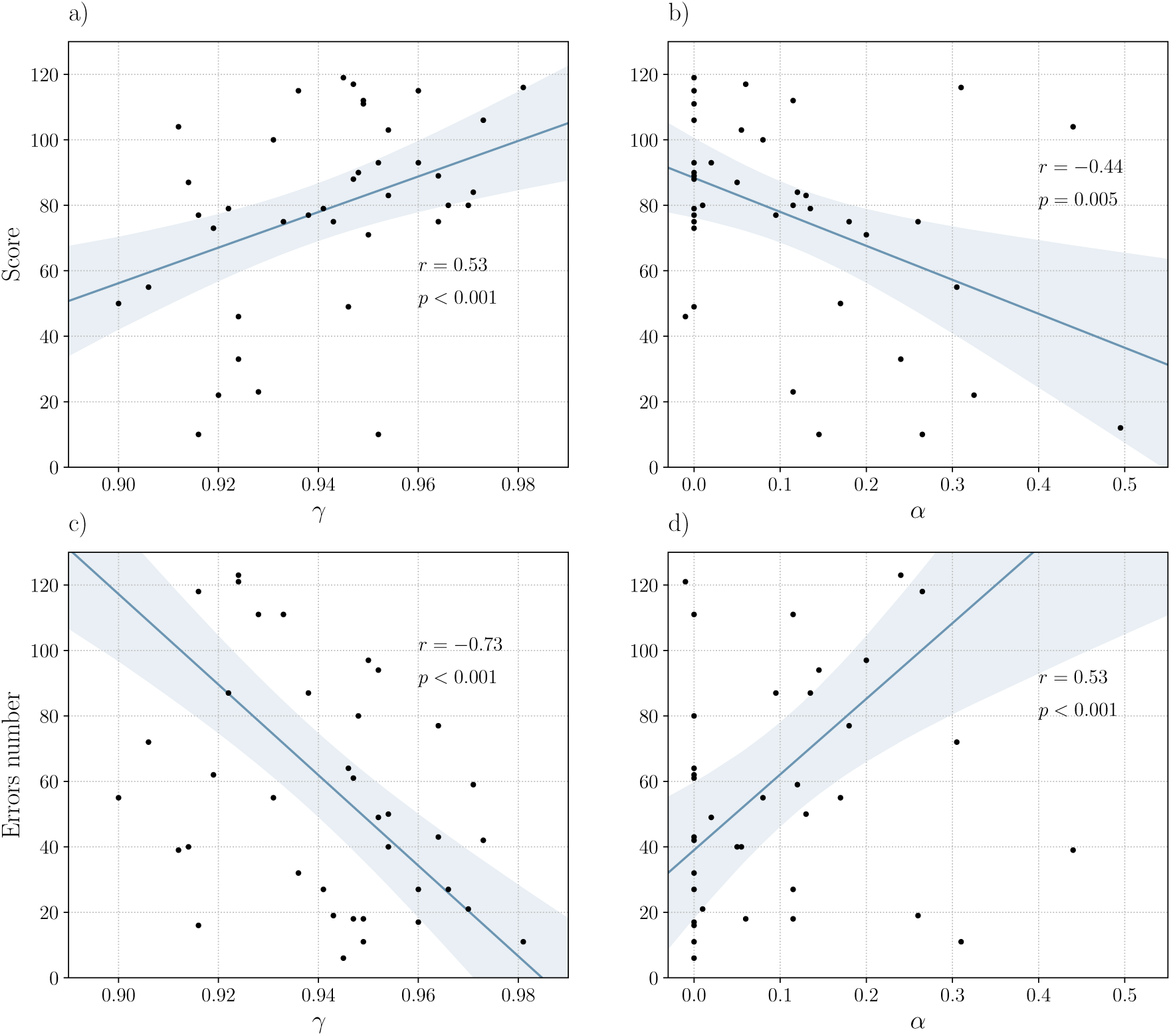
Correlations between participants’ performance and model parameters. **a)** Positive correlation between the farsightedness parameter *γ* and the participants’ scores. Higher values of *γ* are associated with better performance, with Pearson *r* = 0.53 and *p <* 0.001. **b)** Nega-tive correlation between the affordance coefficient *α* and performance, indicating that participants who weighed affordances more heavily tended to score lower, with Pearson *r* = 0.44 and *p* = 0.005. The shaded area represents the 95% confidence interval. **c)** Negative correlation between the far-sightedness parameter *γ* and the number of errors made by participants, with higher values of *γ* associated with fewer errors. Pearson *r* = 0.73 and *p <* 0.001. **d)** Positive correlation between the affordance coefficient *α* and the number of errors. Participants who considered affordances more tended to make more errors. Pearson *r* = 0.53 and *p <* 0.001.

Furthermore, we examined the number of errors made by each participant throughout the experiment, where an error was defined as falling into the water during a jump. Our results reveal a negative correlation between the farsightedness parameter *γ* and the number of errors, suggesting that more farsighted participants tended to make fewer errors (Figure 7.c). Furthermore, the results reveal a positive correlation between affordance coefficients *α* and number of errors, potentially reflecting the fact that participants who made more (fewer) errors tended to assign more (less) importance to the relative difficulty of making safe or risky jumps (Figure 7.c).

To further investigate the role of affordances in jump performance, we examined the relationship between *α* and jump success probability. For consistency with the selection criteria used in other analyses and our definition of decision points, we only focused on jumps starting from large platforms. Furthermore, we only included in the analyses those failed jumps where it was clear which rock the participant was aiming to reach; instead, we discarded ambiguous jumps in which participants landed in the water near their starting rock (Figure 8). These ambiguous cases accounted for approximately 65.5% of all failed jumps (1693 out of 2583), primarily consisting of jumps initiated too late or cases where participants briefly touched the water after landing close to platform edges.

**Figure 8:**
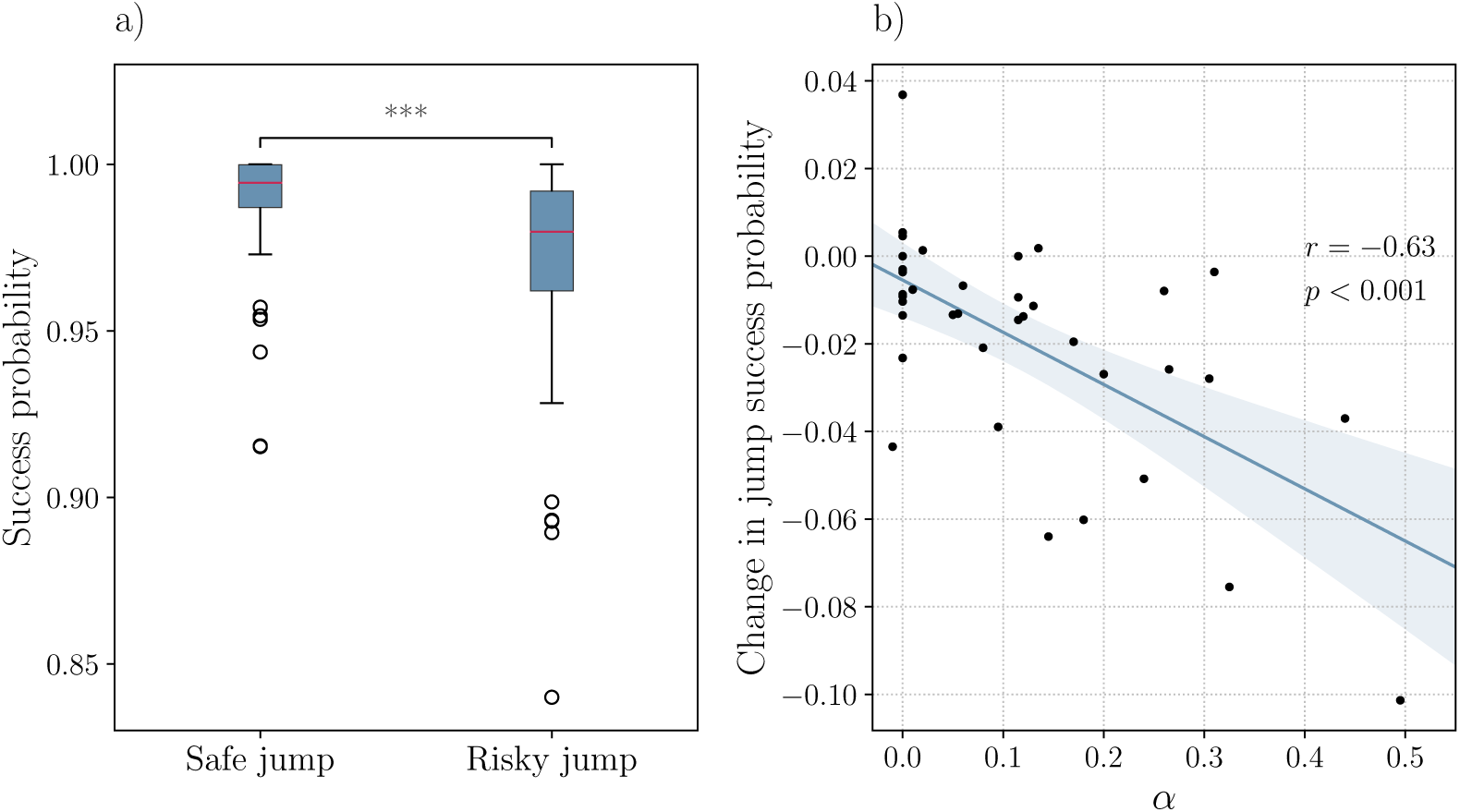
Success probability and its relationship with jump type (safe or risky) and the affordance parameter. **a)** Success probability of safe versus risky jumps, starting from large platforms. Success rates are significantly lower for risky jumps, indicating their increased difficulty. **b)** Correlation between the affordance parameter *α* and the change in success probability when making risky jumps. Participants with lower *α* values show minimal changes in success probability from safe to risky jumps, suggesting a lower sensitivity to affordances, while higher *α* participants exhibit a stronger differentiation between the two types of jumps.

Our analysis reveals that success rates were significantly smaller with risky compared to safe jumps, highlighting the increased difficulty of the former (Figure 8.a). Furthermore, our analysis reveals a negative correlation between *α* and the change in success probability from safe to risky jumps (Figure 8.b). Specifically, participants with lower *α* values show minimal changes in success probability, suggesting that affordance-related factors may not be relevant to them, as all jumps, whether safe or risky, are equally easy for them. In contrast, higher *α* participants are more sensitive to the relative difficulty posed by risky jumps.

Finally, we compared the Bayesian Information Criterion (BIC) across a family of nested models (Table 7). These include one model with no free parameters (the *shortest-path agent*, with *α* = 0 and *γ* = 1), three models with one free parameter (the *utility-only agent*, with *α* = 0 and free *γ*; the *affordance-only agent*, with free *α* and *γ* = 0; and the *no-discount agent*, with free *α* and *γ* = 1), and one agent with two free parameters (the *full-EV agent*, with free *α* and *γ*). Each model captures different combinations of farsightedness and sensitivity to jump difficulty. The full-EV agent achieves the lowest BIC, indicating that participants’ decisions are best captured by a model that integrates both local jump difficulty and long-term path efficiency.

**Table 7:** Model comparison. Mean and standard deviation of log-likelihood and BIC (Bayesian Information Criterion) are reported across participants for five models with increasing complexity. The *shortest-path agent* (*γ* = 1, *α* = 0) has no free parameters. The *utility-only agent* (free *γ*, *α* = 0), *affordance-only agent* (*γ* = 0, free *α*), and *no-discount agent* (*γ* = 1, free *α*), each have one free parameter per participant. The *full-EV agent* (free *γ* and *α*) has two parameters per participant. Lower BIC values indicate better model fit to participant behavior.

| Model |  | log-likelihood |  | BIC |  |
| --- | --- | --- | --- | --- | --- |
|  |  | Mean | Std | Mean | Std |
| Shortest-path agent | ($\alpha = 0, \gamma = 1$) | -210.0 | 83.1 | 419.9 | 166.2 |
| Utility-only agent | ($\alpha = 0$, free $\gamma$) | -156.3 | 38.6 | 319.2 | 77.2 |
| Affordance-only agent | (free $\alpha$, $\gamma = 0$) | -232.5 | 20.8 | 471.6 | 41.6 |
| No-discount agent | (free $\alpha$, $\gamma = 1$) | -199.2 | 66.4 | 405.0 | 132.7 |
| Full-EV agent |  | **-152.5** | 31.1 | **318.2** | 62.3 |

### 3.6 Learning effects across trials

To investigate whether participants’ behavior changed across the experiment, we examined how performance and timing variables evolved over trials. As shown in Figure 9, the average score increased steadily (panel a), while both preplanning time (panel b) and decision time (panel c) decreased, particularly during the early trials. A linear mixed-effects model with trial number as a fixed effect confirmed that all three trends were statistically significant (*p <* 0.001), indicating a general improvement in task execution over time.

**Figure 9:**
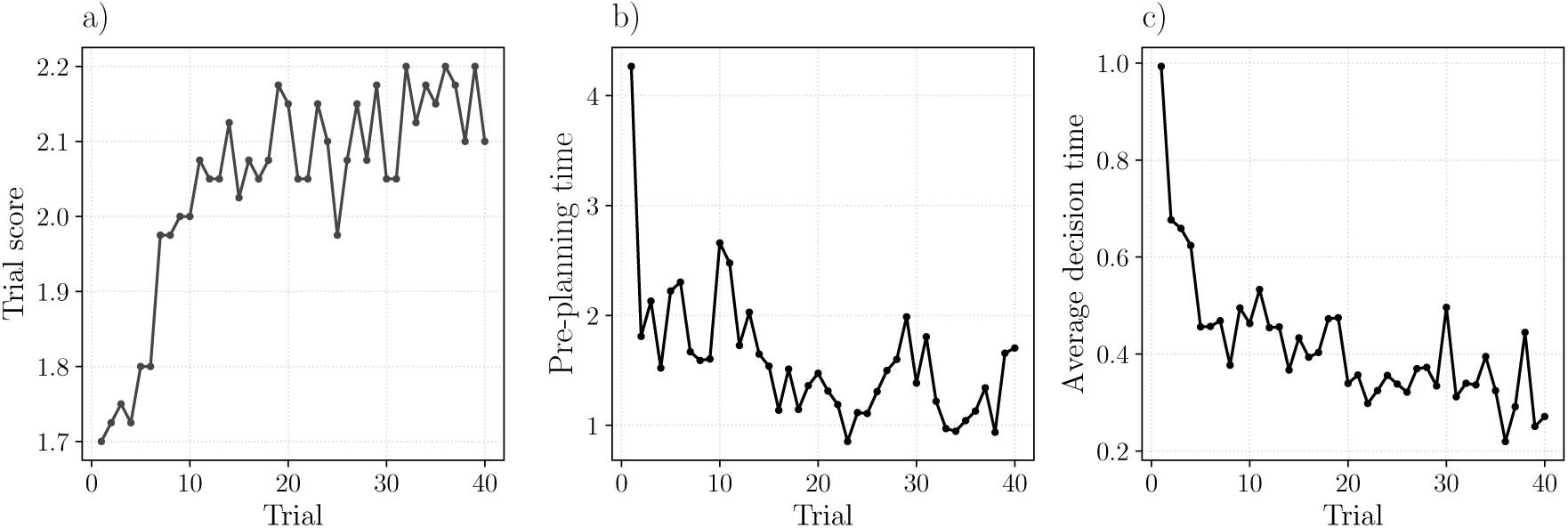
Practice effects over trials. **a)** Average score per trial, showing improvement especially in early trials. **b)** Time spent standing still at the beginning of each level (preplanning time). **c)** Average time spent at decision points within a trial (decision time).

To test whether this learning affected model parameters, we fitted the model separately on the first and second half of the experiment. The two fitted parameters, *γ* and *α*, showed no significant differences (Mann-Whitney U-test, *p* = 0.2 for *γ*, *p* = 0.47 for *α*), justifying our choice to treat all trials together in the model-based analyses of Section 3.5. This result indicates that despite participants’ performance improving across trials, their farsightedness (captured by *γ*) and the weight they assign to affordances (captured by *α*) do not, suggesting no change of strategy.

## 4 Discussion

Every day, we face a myriad of embodied decisions, such as when deciding what path to take in a busy road or which objects to use and in which order to set up a table for breakfast or prepare a backpack for a trekking. These and other tasks present sequences of choices between immediate and more distal affordances and often require simultaneously making sequential decisions and actions (1; 2; 3; 4; 5; 6; 7; 8; 19). Despite their importance, embodied decisions remain poorly investigated. Most studies of decision-making and planning in human psychology and (neuro)economics focus on non-embodied settings and the few studies of embodied decisions tend to focus on single embodied decisions, not sequential decisions (1; 2; 3; 4; 5; 6; 18). Hence, we still do not know whether, when making sequences of embodied decisions, people plan ahead by considering how different choices will lead to new affordances, whether they preplan an entire task before starting or continue planning even while acting, and whether embodied aspects of action execution influence the decisions that are made.

We addressed the above questions by designing a game-like experiment in which participants controlled an avatar required to “cross rivers”, by jumping on a series of rocks of different sizes forming different paths to the goal destination. Crucially, the task presented participants with a series of choices between “safe” and “risky” jumps (corresponding to bigger-and-closer versus smaller-and-farther rocks) and between longer and shorter paths to the goal. This setup permitted us to study how people balance affordance and utility in sequential embodied settings. To get more insights about participants’ strategies, we also developed a computational model implementing jumping decisions as a function of current affordances (“safe” and “risky” jumps) and the utility (length) of the paths that they open up. By fitting the models’ parameters (the weight of the affordance term and the degree of forward planning) to participants’ behavior, we aimed to formalize their decision strategies.

Our behavioral and computational analyses highlight six main findings. First, participants plan ahead their path to the goal rather than simply focusing on the most immediate jumping affordances. At decision points between safe and risky jumps, they balance jumping affordances and their long-term utility, indexed by the length of the paths that they open up and the angle that they form with the goal. Interestingly, despite the angle to goal being a more perceptually available proxy for path utility (albeit potentially misleading), participants take into more consideration the path length, which is a veridical index of utility. The fact that participants show effective planning strategies during “crossing the river” illustrates the effectiveness of embodied tasks to study higher cognitive processes (43).

Second, spatial and embodied components of the task influence participants’ decision strategies. This is possible since, despite being merely virtually embodied, our game requires detailed sensori-motor control of the avatar and the consideration of spatial factors and environmental affordances. Furthermore, the movements of the avatar follow coherent physical and spatial rules and the ease of executing a jump while continuing along the same trajectory (without the motor cost required to change direction) may impact the participant’s decisions. The preference for jumping on rocks in the current direction of movement is stronger in the main experiment, which requires a continuous flow of movement (Experiment 1), is attenuated when we break the flow but still allow participants to rotate in place (Experiment 2) and disappears—but does not reverse—when we “flip” the direction of movement after each jump (Experiment 3). Neither motor cost nor time required to rotate can explain these results if considered in isolation—as participants do not reverse their preference to favor the new direction when rotated to it (Experiment 3), nor do they always select the jump associated with smaller motor cost. This pattern of result suggests instead that participants have a preference for forming and sticking to spatially coherent plans as if their physical momentum plays a modulatory role. Our results add to a growing literature showing that in (virtually) embodied settings, spatial and embodied aspects of the task, such as direction of movement and action dynamics, can influence decisions—and how we assign value and preference to choice offers (10; 21; 14; 15). However, our findings extend previous results in important ways. Previous studies presented choice alternatives (e.g., new targets to be tracked in (10)) abruptly, while participants were acting, but without giving them the possibility to plan multiple steps ahead. By contrast, in our study, participants could see the whole map and plan multiple jumps ahead at the beginning of each trial, before starting movement. Hence, they could have potentially pre-planned a series of decisions not sensitive to action dynamics. Yet, comparing the main experiment (Experiment 1) with a condition in which physical momentum is removed (Experiment 2) reveals that the latter still plays a modulatory role, indicating that participants continue integrating contributions from the unfolding action into their decision process even after having formed a plan (see also below for a discussion of simultaneous planning and action).

Third, participants preplan part of the path at the beginning of the task, before making the first jump. This is evidenced by the fact that preplanning time is influenced by the perceived angle between the first risky jump and the goal and that longer preplanning time can lead to improved solutions (shorter paths). This finding parallels studies of preplanning during more abstract problem solving, such as the fact that humans think for different durations in different contexts (44), highlighting that participants are sensitive to the sequential planning nature of “crossing the river”.

Fourth, planning is not completed before movement but continues during the task. This is shown by the fact that participants’ deliberation time (indexed by slowing down during the task, not related to motor requirements) increases at decision points and even before them. This finding illustrates that in embodied settings, participants can simultaneously plan and act – and that a strict separation between the two processes does not allow accounting for subtle aspects of naturalistic behavior, such as the fact that movement kinematics index decision variables such as decision uncertainty and confidence (45; 46; 47) and that participants can change their mind along the way (48).

Fifth, computational modeling shows that farsighted participants (which assigned greater weight to the utility of future jumps) show better task performance, above and beyond their jumping skill (which varies across participants independent of decision strategy). This finding highlights the possibility to adopt computational models that formalize the usual trade-offs that characterize value-based decisions, such as between the probabilities of the different outcomes (here, of successful jumps) and their utilities (here, path length) but in embodied settings (43). Besides providing a way to quantify participants’ strategies, the variables inferred by the model – namely, affordance and utility landscapes at decision points – could be potentially used a predictors of neural rep-resentations in future studies that address the neuronal underpinnings of embodied decision and planning processes (49; 50; 51; 52).

Finally, we found that participants’ performance increases during the experiment, as reflected by greater score per trial and lower preplanning and decision time. However, the model-based parameters reflecting farsighteness and the weight assigned to affordances remain stable across trials, suggesting that participants do not significantly change their overall decision strategy during the experiment.

Taken together, our results show that when solving an embodied task (even a “virtually embodied” one) – crossing the river by continuously selecting and executing “risky” or “safe” jumps – participants adopt effective planning strategies that balance immediate jump affordances and long-term path utility. Notably, participants’ decision strategies are influenced by embodied characteristics of the task such as their current travel direction and reciprocally, their movement speed is influenced by the necessity to continue planning along the way. These findings emphasize the interplay between decisions and actions in embodied tasks, shedding light on the cognitive mechanisms underlying real-world planning-while-acting scenarios. Furthermore, they underscore the importance of studying higher level processes such as planning in settings that reflect the ecological conditions of our everyday decisions and actions.

This study has various limitations that will need to be addressed in future studies. First, we asked participants to make sequences of jumping decisions and we analysed participants’ choices at both the first and the subsequent decision points – but the two analyses pose different challenges. The results at the first decision point are more straightforward to interpret, since they directly relate to our main experimental manipulation. However, the interpretation of the results at subsequent decision points is complicated by the fact that some factors that we considered (e.g., travel and goal direction, path length) can be sometimes correlated – which is something to be expected in sequential tasks in ecologically valid settings. The coherence of our results across the first and the subsequent decision points is reassuring and suggests that possible confounds played a minor role. Further strengthening this interpretation is the fact that our model comparison reveals that the GLMM without interaction between the two factors of our design best explains the data on subsequent decision points. This suggests that participants might treat these two factors separately, making confounds between them less likely. It is also important to remark that our approach using both linear (GLMM) and computational (EV) modeling further mitigates the concerns, addressing complementary aspects of the decision setup. For example, while at the first decision point the gain of selecting the shortest path is always 2 jumps, this is not always the case across the subsequent decision points. Consider for instance that a risky path intersecting a safe path oriented more or less directly towards the goal could create decision points where the difference in path length between the two options is greater or smaller. While this fact is not considered in the GLMM, the EV model takes it into account, since the difference in EV between two options is proportional to their different path length. Summing up, there is always a tension between controlling every factor precisely at the cost of simplifying the decision setup (as done in many laboratory studies) and addressing the richness of ecological scenarios, at the cost of having additional factors. Our approach in this study provides a compromise – as it maps classical factors of two-alternative forced choices considered in neuroeconomics, like probabilities and utilities, into embodied factors that can be experimentally manipulated and formalized with the aid of computational models. Yet, future studies might devise additional ways to disentangle more clearly the factors of interest ideally without losing the richness of ecological settings.

Another limitation of our study is that it only addresses some of the key factors that characterize embodied choices. Future designs could address additional embodied choice factors, such as different jumping costs (which in our design does not depend on rock distance) and the need to explore the environment (which in our design is largely sidestepped). Similarly, future studies might manipulate the distance between rocks and their size independently and include additional affordance-related factors (e.g., slippery rocks) – in order to study how participants integrate these different affordances when making jumping decisions. Furthermore, while in this study we have identified some key factors that influence participants’ decisions, additional studies are needed to identify the cognitive processing underlying their decision dynamics. For example, we interpreted the additional time participants spent at decision and pre-decision points compared to non-decision points as “planning time”. However, alternative explanations are possible, such as recalling the plan or checking if the orientation of the avatar is correct. Designing future experiments and richer computational models could help identify more clearly the cognitive processing underlying sequential embodied decisions. Finally, future studies might replicate and extend these findings during real-world jumping and river crossing tasks, measure additional variables, such as full body kinematics, eye movements and physiological parameters, and exploit more advanced computational models, in order to shed more light on how people address sequential embodied tasks “in the wild” (53; 54; 55).

Summing up, our study illustrates a rich interplay between action and cognition in real-world planning-while-acting scenarios, revealing that people effectively balance immediate affordances and long-term utilities of alternative action plans, plan ahead before the task but continue deliberating during it, and consider geometric, spatial, and embodied components of the task during decision-making. From a methodological point of view, studying sequential choices in embodied settings reveals subtle aspects of decision processes that might be relevant to understand everyday cognition, but might be missed when only focusing on non-embodied settings that remove spatial and spatial factors from the choice, as usually done in (neuro)economics. Our study provides a demonstration of how concepts from (neuro)economics can be merged with concepts on embodiment into a richer picture that’s missing from both of those approaches.

## Acknowledgements

The authors wish to thank Thomas Thiery for helping to develop the original concept of the river crossing task. This research received funding from the European Research Council under the Grant Agreement No. 820213 (ThinkAhead), the Italian National Recovery and Resilience Plan (NRRP), M4C2, funded by the European Union – NextGenerationEU (Project IR0000011, CUP B51E22000150006, “EBRAINS-Italy”; Project PE0000013, “FAIR”; Project PE0000006, “MNESYS”), and the Ministry of University and Research, PRIN PNRR P20224FESY and PRIN 20229Z7M8N. PC was supported by a grant from the Natural Sciences and Engineering Research Council of Canada (RGPIN-2022-05345). The funders had no role in study design, data collection and analysis, decision to publish, or preparation of the manuscript. The GEFORCE Quadro RTX6000 and Titan GPU cards used for this research were donated by the NVIDIA Corporation.

## A Supplementary materials

### A.1 Current travel direction influences subsequent jump choices: Supplementary analysis

This section complements the analysis of how current travel direction influences jump choices of Section 3.2.

Given that each decision point presents two possible choices – either a *risky* or a *safe* jump – we analyzed these two possibilities separately. First, we considered how often participants selected the *risky* jump, based on the trajectory angle. For ease of analysis, we categorized the trajectory angles (which are continuous variables) into a categorical variable with three levels: i) 0^◦^ to 45^◦^; ii) 45^◦^ to 90^◦^; iii) greater than 90^◦^. This classification reflects the arrangement of rocks in a hexagonal grid, where typical angles between directions (ignoring jitter) are around 0^◦^, 60^◦^, and occasionally 120^◦^ or 180^◦^ when participants backtrack (note that the latter condition occurs rarely in our dataset, about 1% of the jumps). The binning procedure was designed to ensure equal-sized angular ranges across conditions. Note that trajectory angles span both negative and positive values (e.g., from –180° to +180°), depending on the direction of deviation relative to the current travel vector. The interval centered on 0° covers [–45°, 45°], while the interval centered on 60°, spans both [45°, 90°] and [–90°, –45°], preserving symmetry and consistent bin width. The final interval includes all remaining angles greater than 90° or less than –90° (i.e., from 90° to 180° and –180° to –90°). Although this range is broader, we chose not to subdivide it further due to the low number of data points in this category, which would limit the reliability of statistical comparisons. We fit a GLMM to model the probability of choosing a *risky* jump as a function of the trajectory angle (Table S1). Using the [0^◦^, 45^◦^] class as the baseline, both other angle classes exhibit significant differences (*p <* 0.001). This indicates that the trajectory angle significantly influences participants’ *risky* jump decisions. The results are further illustrated in Figure S1.b, showing that all conditions are significantly different from each other (see Table S6 for the post-hoc comparisons results).

We next made the same analysis for *safe* jumps (Figure S1.c, Table S2). Similar to *risky* jumps, the results reveal a significant effect of the trajectory angle (*p <* 0.001) and that the probability of performing the *safe* jump decreases significantly as the trajectory angle increases (see Table S7 for post-hoc comparison results).

These results align with those reported in the main text, showing that participants are more likely to select either jump option, when their current travel direction is aligned with it. This suggests that participants do not consider only the static costs and benefits of the choice alternatives but also the continuity of movement and the ease with which they can execute an action within the context of their current dynamic state, demonstrating the relevance of the current direction of travel for the choice. This effect emerges in our task since the sequential choices are not independent from one another and the task does not break down the natural contingency between movement and subsequent perception. This feature is usually absent from non-embodied, sequential decisions often addressed in (neuro)economics, which comprise series of discrete and independent choices (56; 43).

**Figure S1:**
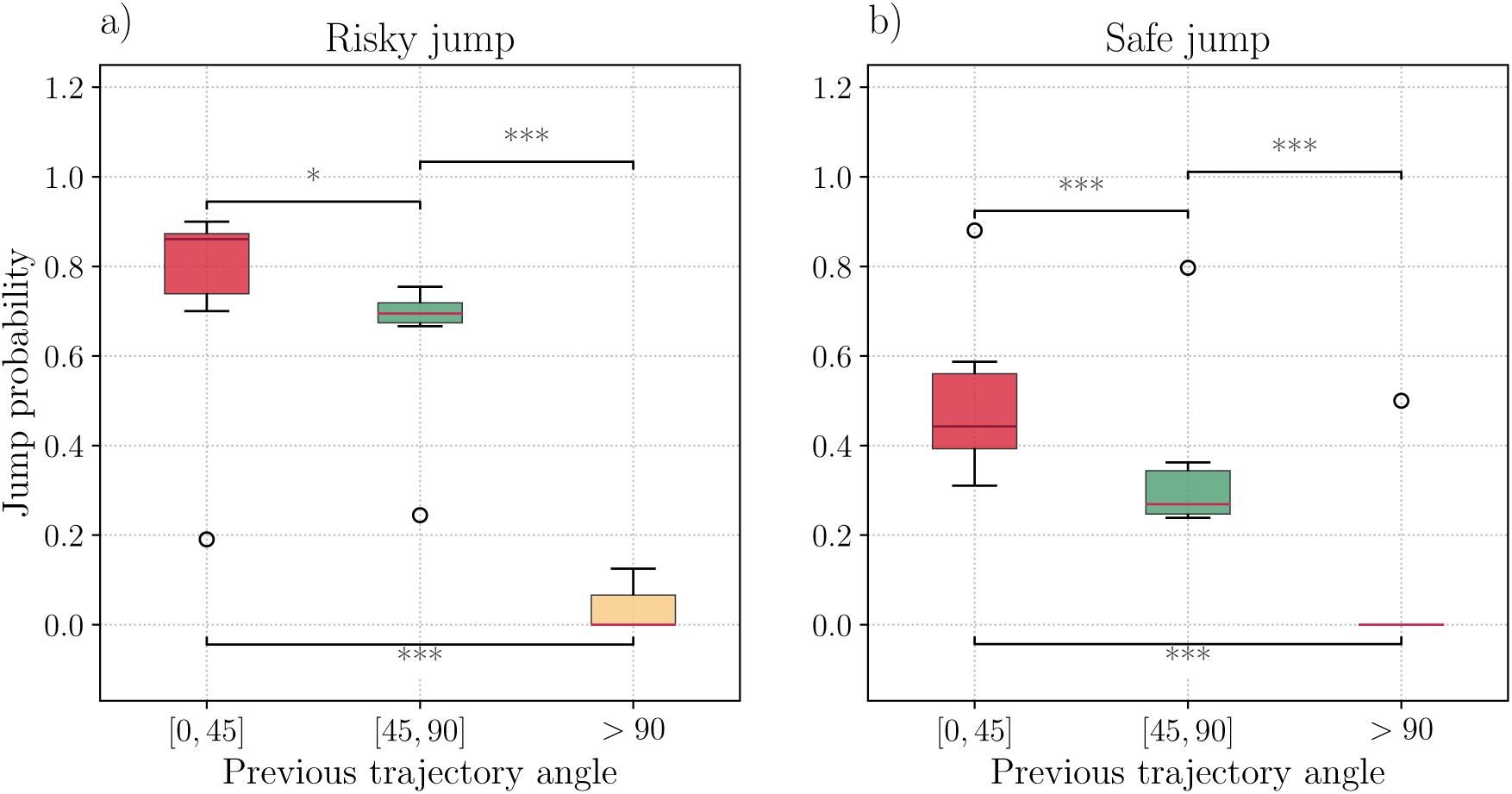
Influence of trajectory angle on *risky*and *safe* jump probabilities. **a)** Boxplot showing the probability of performing a *risky* jump as a function of the previous trajectory angle. The trajectory angle was divided into three categories: [0^◦^, 45^◦^], [45^◦^, 90^◦^], and greater than 90^◦^. Significant differences are observed across all three categories (*p <* 0.001) indicating a sharp decrease in *risky* jump probability as the angle increases. **b)** Boxplot showing the probability of performing a *safe* jump, analyzed in the same way as the *risky* jumps. Here, too, the probability decreases significantly as the trajectory angle increases, demonstrating the same trend observed for *risky* jumps.

**Table S1:** GLMM results for *risky* jump probabilities as a function of trajectory angle. The trajectory angle is divided into three categories: [0^◦^, 45^◦^], [45^◦^, 90^◦^], and greater than 90^◦^. The [0^◦^, 45^◦^] category serves as the baseline. The table shows significant decreases in *risky* jump probability as the trajectory angle increases. Random effects account for individual participant variability.

|  | Fixed effects |  |  |  |  | Random effects |
| --- | --- | --- | --- | --- | --- | --- |
|  | Estimate | Std Error | [0.025 | 0.095] | p-value | Variance |
| Intercept | -0.456 | 0.106 | -0.664 | -0.247 | < **0.001** | 0.265 |
| Angle $[45^\circ, 90^\circ]$ | -3.022 | 0.142 | -3.301 | -2.744 | < **0.001** | |
| Angle $> 90^\circ$ | -1.013 | 0.089 | -1.187 | -0.839 | < **0.001** | |

**Table S2:** GLMM results for *safe* jump probabilities as a function of trajectory angle. The trajectory angle is divided into three categories: [0^◦^, 45^◦^], [45^◦^, 90^◦^], and greater than 90^◦^. The [0^◦^, 45^◦^] category serves as the baseline. The table shows significant decreases in *safe* jump probability as the trajectory angle increases. Random effects account for individual participant variability.

|  | Fixed effects |  |  |  |  | Random effects |
| --- | --- | --- | --- | --- | --- | --- |
|  | Estimate | Std Error | [0.025 | 0.095] | p-value | Variance |
| Intercept | -1.079 | 0.119 | -1.313 | -0.846 | < **0.001** | 0.318 |
| Angle $[45^\circ, 90^\circ]$ | -2.369 | 0.164 | -2.691 | -2.048 | < **0.001** | |
| Angle $> 90^\circ$ | -0.559 | 0.101 | -0.756 | -0.361 | < **0.001** | |

### A.2 Post-hoc comparison tables for all LMM, GLM and GLMM analysis

Each table provides the estimated effects, standard errors, confidence intervals, and p-values for each contrast. Significant results are highlighted. All *p*-values have been corrected with Tukey’s Honestly Significant Difference (HSD) test.

**Table S3:** Pairwise post-hoc comparisons for the probability of making a *risky* jump at the first decision point. The contrasts between different combinations of path length and angle are shown. For brevity, larger/smaller refer to the angle to the goal, and longer/shorter refer to the path length.

| Contrast | Estimate | Std Error | [0.025 | 0.095] | p-value |
| --- | --- | --- | --- | --- | --- |
| Larger / Longer - Smaller / Longer | -0.721 | 0.161 | -1.135 | -0.307 | < **0.001** |
| Larger / Longer - Larger / Shorter | -1.082 | 0.158 | -1.487 | -0.677 | < **0.001** |
| Larger / Longer - Smaller / Shorter | -2.655 | 0.190 | -3.142 | -2.168 | < **0.001** |
| Smaller / Longer - Larger / Shorter | -0.361 | 0.157 | -0.765 | 0.044 | 0.100 |
| Smaller / Longer - Smaller / Shorter | -1.934 | 0.188 | -2.416 | -1.452 | < **0.001** |
| Larger / Shorter - Smaller / Shorter | -1.573 | 0.182 | -2.042 | -1.104 | < **0.001** |

**Table S4:**
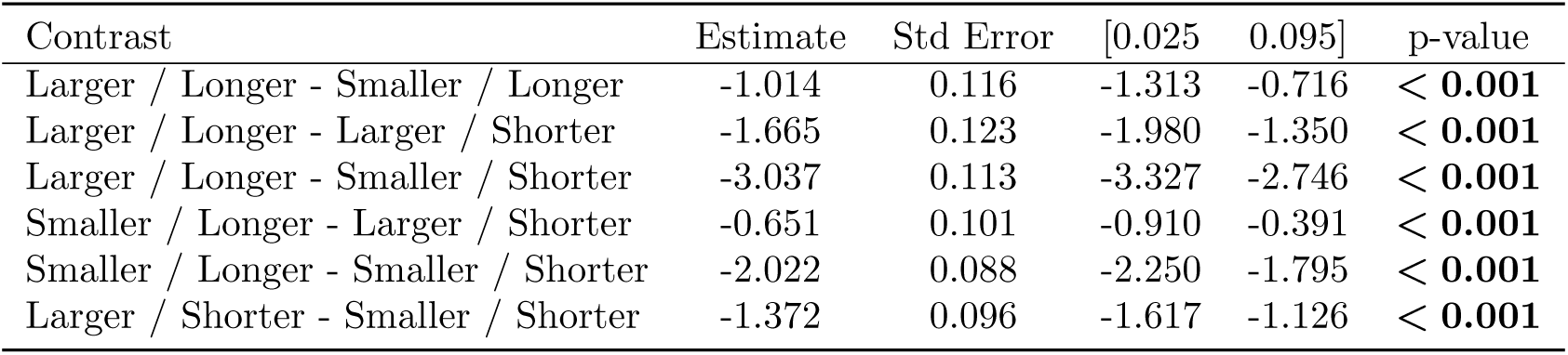
Pairwise post-hoc comparisons for the probability of making a *risky* jump across all decision points. The contrasts represent combinations of angle to the goal (larger/smaller) and path length (longer/shorter).

**Table S5:** Pairwise post-hoc comparisons for the pre-planning time. The contrasts represent combinations of angle to the goal (larger/smaller) and path length (longer/shorter).

| Contrast | Estimate | Std Error | [0.025 | 0.095] | p-value |
| --- | --- | --- | --- | --- | --- |
| Larger / Longer - Smaller / Longer | 0.171 | 0.052 | 0.038 | 0.304 | **0.005** |
| Larger / Longer - Larger / Shorter | -0.061 | 0.052 | -0.194 | 0.072 | 0.638 |
| Larger / Longer - Smaller / Shorter | 0.110 | 0.073 | -0.079 | 0.298 | 0.438 |
| Smaller / Longer - Larger / Shorter | -0.232 | 0.073 | -0.420 | -0.044 | **0.008** |
| Smaller / Longer - Smaller / Shorter | -0.061 | 0.052 | -0.194 | 0.072 | 0.638 |
| Larger / Shorter - Smaller / Shorter | 0.171 | 0.052 | 0.038 | 0.304 | **0.005** |

**Table S6:** Post-hoc comparison of trajectory angle categories for *risky* jump probabilities. Results of the post-hoc pairwise comparisons between the three trajectory angle categories ([0^◦^, 45^◦^], [45^◦^, 90^◦^], and greater than 90^◦^) for *risky* jump probabilities.

| Contrast | Estimate | Std Error | [0.025 | 0.095] | p-value |
| --- | --- | --- | --- | --- | --- |
| $[0^\circ, 45^\circ] - [45^\circ, 90^\circ]$ | 0.896 | 0.074 | 0.722 | 1.070 | $< \mathbf{0.001}$ |
| $[0^\circ, 45^\circ] - \text{Greater than } 90^\circ$ | 4.274 | 0.201 | 3.803 | 4.745 | $< \mathbf{0.001}$ |
| $[45^\circ, 90^\circ] - \text{Greater than } 90^\circ$ | 3.378 | 0.196 | 2.919 | 3.837 | $< \mathbf{0.001}$ |

**Table S7:**
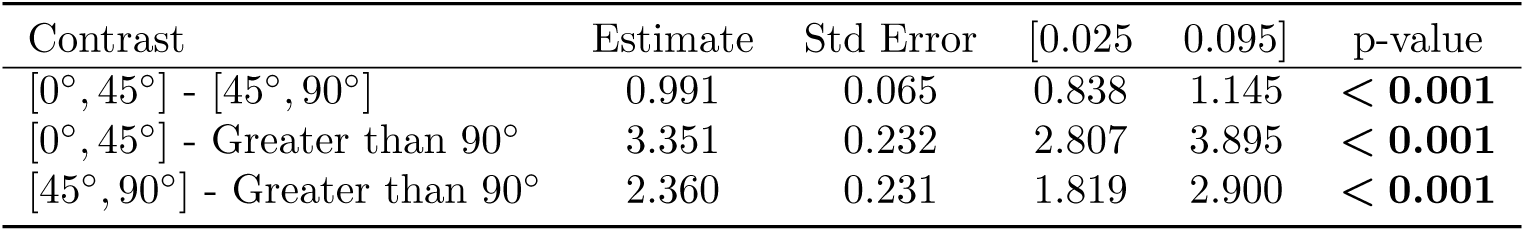
Post-hoc comparison of trajectory angle categories for *safe* jump probabilities. Results of the post-hoc pairwise comparisons between the three trajectory angle categories ([0^◦^, 45^◦^], [45^◦^, 90^◦^], and greater than 90^◦^) for *safe* jump probabilities.

| Contrast | Estimate | Std Error | [0.025 | 0.095] | p-value |
| --- | --- | --- | --- | --- | --- |
| $[0^\circ, 45^\circ] - [45^\circ, 90^\circ]$ | 0.991 | 0.065 | 0.838 | 1.145 | $< \mathbf{0.001}$ |
| $[0^\circ, 45^\circ] - \text{Greater than } 90^\circ$ | 3.351 | 0.232 | 2.807 | 3.895 | $< \mathbf{0.001}$ |
| $[45^\circ, 90^\circ] - \text{Greater than } 90^\circ$ | 2.360 | 0.231 | 1.819 | 2.900 | $< \mathbf{0.001}$ |

**Table S8:** Post-hoc comparison of plannin time as a function of platform type. The contrasts represent the possible combinations of platform types.

| Contrast | Estimate | Std Error | [0.025 | 0.095] | p-value |
| --- | --- | --- | --- | --- | --- |
| Nondecision - Predecision | -0.049 | 0.010 | -0.073 | -0.025 | $< \mathbf{0.001}$ |
| Nondecision - Decision | -0.114 | 0.008 | -0.134 | -0.095 | $< \mathbf{0.001}$ |
| Predecision - Decision | -0.065 | 0.012 | -0.094 | -0.037 | $< \mathbf{0.001}$ |

**Table S9:** Post-hoc comparison of jump choice as a function of experiment setup and change in direction. The contrasts represent the possible combinations of experiment (1 to 3) and direction of the new jump (current direction or new direction).

| Contrast | Estimate | Std Error | [0.025 | 0.095] | p-value |
| --- | --- | --- | --- | --- | --- |
| Current Dir / Exp 1 - New Dir / Exp 1 | 1.282 | 0.049 | 1.144 | 1.421 | $< \mathbf{0.001}$ |
| Current Dir / Exp 1 - Current Dir / Exp 2 | 0.296 | 0.053 | 0.144 | 0.447 | $< \mathbf{0.001}$ |
| Current Dir / Exp 1 - Current Dir / Exp 3 | 0.640 | 0.052 | 0.492 | 0.789 | $< \mathbf{0.001}$ |
| New Dir / Exp 1 - New Dir / Exp 2 | -0.296 | 0.053 | -0.447 | -0.144 | $< \mathbf{0.001}$ |
| New Dir / Exp 1 - New Dir / Exp 3 | -0.640 | 0.052 | -0.789 | -0.492 | $< \mathbf{0.001}$ |
| Current Dir / Exp 2 - New Dir / Exp 2 | 0.691 | 0.057 | 0.528 | 0.855 | $< \mathbf{0.001}$ |
| Current Dir / Exp 2 - Current Dir / Exp 3 | 0.345 | 0.056 | 0.184 | 0.506 | $< \mathbf{0.001}$ |
| New Dir / Exp 2 - New Dir / Exp 3 | -0.345 | 0.056 | -0.506 | -0.184 | $< \mathbf{0.001}$ |
| Current Dir / Exp 3 - New Dir / Exp 3 | 0.002 | 0.055 | -0.156 | 0.160 | 1.000 |

